# CIP2A tetramerization is required for mitotic DNA repair

**DOI:** 10.64898/2026.08.22.746418

**Authors:** Lauren de Haan, Rebecca Schneeweiss, Marco Barazas, Milo L. Kaptein, Femke J. Bakker, Anna Dekker, Ekaterina Ovcharenko, Soraya S. Beneddine, Angela Graf, Dimitra Paouneskou, Bert van de Kooij, Raimundo Freire, Verena Jantsch, Jos Jonkers, H. Rudolf de Boer, Marcel Tijsterman, Marcel A. T. M. van Vugt, Pim J. Huis in ‘t Veld

## Abstract

DNA lesions that persist in mitosis threaten genome stability. These lesions recruit TOPBP1-CIP2A, a complex crucial to tether and process damaged DNA on mitotic chromosomes. Importantly, CIP2A is synthetic lethal in *BRCA1/2* mutant cancer cells. However, mechanistic insight into the function of CIP2A is lacking. Here we provide first structural insights into full-length CIP2A and reveal how CIP2A dimers, 46 nm in length, assemble via conserved C-terminal motifs into dumbbell-like tetramers. DNA repair signatures demonstrate that CIP2A tetramerization is crucial for mitotic DNA repair by polymerase theta (POLQ). Tetramerization of CIP2A is needed to recruit POLQ to mitotic DNA lesions and to prevent micronucleation upon replication-born DNA lesions. Tetramerization-defective CIP2A mutants specifically decrease survival of *BRCA1/2* mutant cancer cells in a dominant negative manner. Combined, we demonstrate that CIP2A tetramers are crucial components of a mitotic DNA-tethering and repair scaffold, which processes replication-mediated DNA lesions that persist in mitosis.

## Main

DNA replication is continuously challenged, even under physiological conditions^1^. To resolve replication-mediated DNA lesions, cells have evolved various DNA repair mechanisms, which are coordination with cell-cycle checkpoints to maintain genome integrity^2^. Cancer cells commonly show increased levels of perturbed DNA replication, for instance caused by cancer-associated DNA repair defects. Specifically, mutations in DNA repair genes *BRCA1* or *BRCA2* cause instability of stalled replication forks and incomplete replication^3–6^. The resulting DNA lesions and interconnected DNA molecules can be transmitted into mitosis, where they need to be processed in time to allow faithful segregation of chromosomes^7–11^. Since canonical DNA repair pathways are thought to be largely inactivated during mitosis, cells employ dedicated mitotic DNA processing machineries to achieve this^12–15^. Importantly, CIP2A is essential for survival of cancer cells with elevated levels of mitotic DNA damage, as demonstrated for basal-like breast cancer cells^16^ and, in particular, homologous-recombination (HR)-defective cancer cells^17,18^. Thus, mechanistic insight into how the mitotic response to DNA damage is organized by CIP2A is crucial to capitalize on this therapeutic potential.

CIP2A, initially identified as an inhibitor of the PP2A phosphatase^19^, localizes to the cytoplasm during interphase and only gains access to chromosomes when the nuclear envelop breaks down in mitosis^20^. Together with its mitotic binding partner TOPBP1^17,16^, CIP2A tethers fragmented chromosomes^21,22^. Simultaneously, the CIP2A-TOPBP1 complex is needed to recruit DNA processing factors, including the SMX tri-nuclease complex^23–26^ and the DNA repair polymerase POLQ^24,27,28^, to DNA lesions. In contrast to other components of the mitotic response to DNA lesions for which a molecular framework is beginning to emerge, the function of CIP2A in mitosis remains enigmatic.

## Results

### CIP2A dimers assemble into tetramers

To understand the function of CIP2A, we analyzed its structure. Previous structural information of the CIP2A N-terminal domain^29^ and *in silico* predictions^23^ indicated that CIP2A forms dimers with a globular domain and a coiled coil extension (Fig. 1A). This model predicts a separation of more than 40 nm between two regions of CIP2A that are critical for genome maintenance during mitosis: a conserved surface on the globular domain that binds TOPBP1^17,30^, and a short C-terminal extension with an unknown but essential function^23^ (Fig. 1A). We purified full-length human CIP2A using affinity chromatography and sedimentation on a density gradient (Fig. 1B). Mass photometry showed two CIP2A species with molecular weights of approximately 200 and 400 kDa, corresponding to the expected mass of CIP2A dimers and tetramers (Fig. 1C). The direct visualization of full-length CIP2A using electron microscopy (EM) after platinum-shadowing revealed highly elongated particles (Fig. 1D). A fraction of the observed particles measured 46 nm from end-to-end, corresponding to predicted structures of full-length CIP2A dimers and providing first experimental support for a near-continuous parallel dimeric coiled coil in CIP2A (Fig. 1E). Strikingly, most particles displayed a dumbbell-like appearance and measured 90 nm, approximately twice the predicted length of the dimer, suggesting that dimers of CIP2A assemble via an end-to-end configuration into tetramers. Of note, we observed a higher tetramer: dimer ratio by EM than by mass photometry, consistent with the low concentration (20 nM) used for mass photometry.

**Figure 1:**
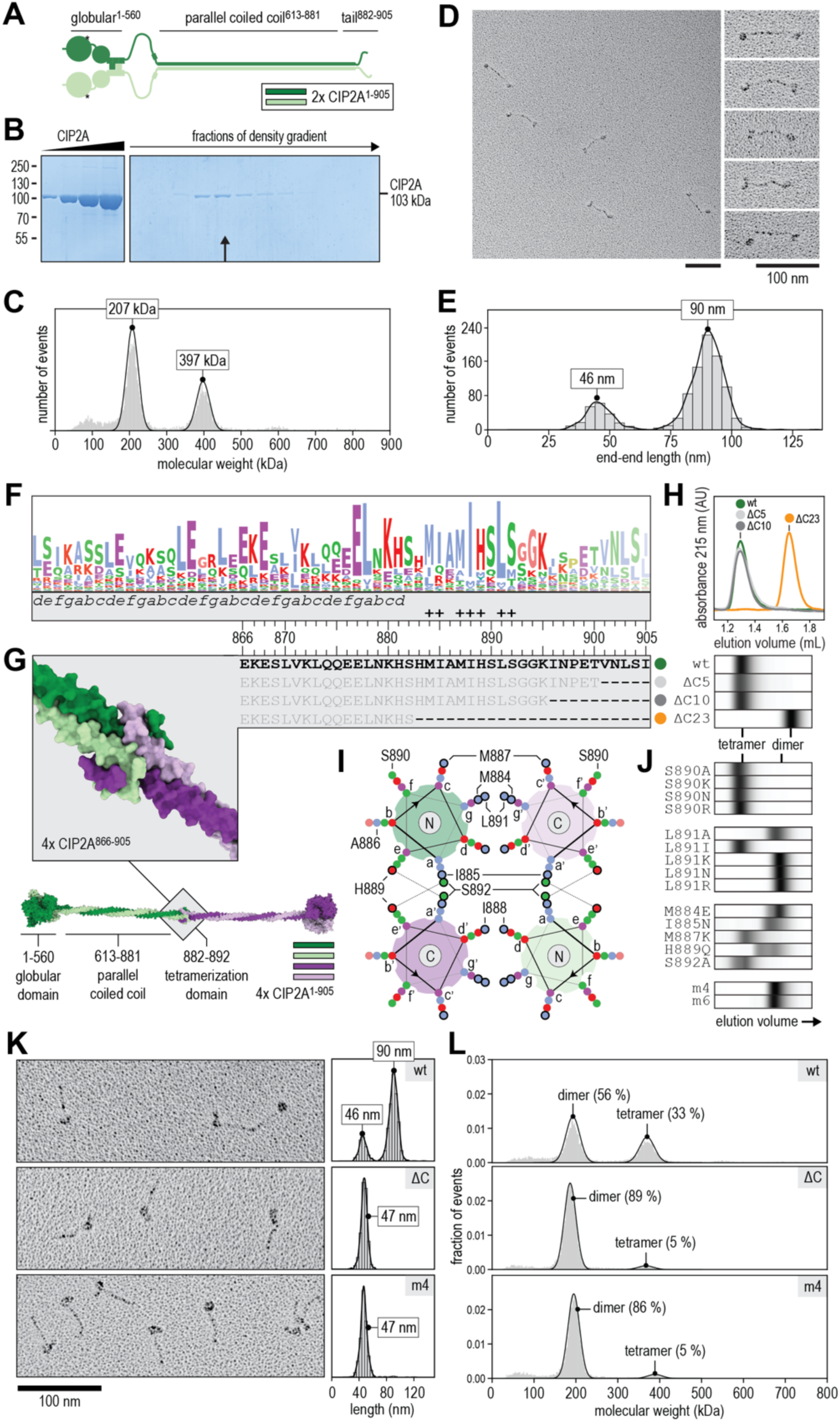
Dimers of CIP2A interact via a C-terminal antiparallel coiled coil. (**A**) Schematic representation of two CIP2A molecules that dimerize via an extended parallel coiled coil and via their globular domains. Asterisks indicate TOPBP1-binding sites. (**B**) Analysis of full-length recombinant human CIP2A by SDS-PAGE and Coomassie staining before (left panel) and after (right panel) sedimentation on a density gradient. CIP2A is 102 kDa in its native form and 103 kDa in its tagged form. The black arrow indicates a fraction used for further analysis. (**C**) Mass photometry of recombinant full-length human CIP2A. Bin size 2.25 kDa, 9,689 events. Black traces show Gaussian kernel density estimation (KDE) to obtain a bin-independent representation of mass distributions in the 100-300 and 300-500 kDa range. (**D**) Electron micrograph of full-length human CIP2A after low-angle rotary metal shadowing. The five particles visible in the left panel are depicted on the right after rotation. (**E**) CIP2A particle lengths (n = 1007) were traced from the beginning of a globular domain to the end. Bin size, 4 nm. Black traces show Gaussian kernel density estimations (KDE) of particles <65 and >65 nm to obtain bin-independent representations of particle length. (**F**) Amino acid frequency for the most C-terminal 60 residues, corresponding to human CIP2A^846–905^, in 401 CIP2A homologs. Side-chain properties are indicated via a Clustal colouring scheme with e.g. hydrophobic (blue), hydrophilic (green), acidic (purple), and basic (red). See Suppl. Fig. 1 for more information. The grey box shows the predicted a-g register of the parallel dimeric coiled coil. The residues that contribute most to CIP2A tetramerization, according to PDBe PISA based on the structure shown in panel G, are marked with a +.(**G**) Structural prediction of four copies of CIP2A^781–905^ with residues 866-905 shown in structure and residue-per-residue (top row, green dot). Residues absent in the struncated constructs (light grey, grey, and orange dots) are indicated with a dash. Predicted structures of CIP2A^1–905^ dimers were superposed with either ends of the predicted CIP2A^781–905^ tetramers to generate the model of 4 copies CIP2A^1–905^. (**H**) Size-exclusion chromatography profiles of CIP2A^753–905^ (green), CIP2A^753–900^ (light grey) CIP2A^753–895^ (grey), and CIP2A^753–882^ (orange) constructs. Retention volumes indicate CIP2A tetramerization (1.2-1.4 mL) and tetramerization (1.6-1.8 ml). Elution profiles were summarized in strips with absorbance values represented as a greyscale. (**I**) Helical-wheel representation of four CIP2A^866–893^ alpha helices that assemble into an antiparallel coiled coil. A cross-section of the predicted structure shown in panel G with consistent colours. Letters a-g indicate the coiled coil register. The N or C in the helical wheel indicate the orientation of the alpha helix. Amino acids are colored as in panel F. The residues that contribute most to CIP2A tetramerization, according to PDBe PISA based on the structure shown in panel G, are outlined in bold. The dashed lines between H889 and S892 indicate a predicted hydrogen-bond. (**J**) Size-exclusion chromatography analysis of CIP2A^753–905^ constructs with all indicated variants summarized as in panel H. (**K**) Representative micrographs and length measurements of CIP2A^wt^, CIP2A^ΔC^ (n = 429), CIP2A^m4^ (n = 607). Length distribution of CIP2A^wt^ is also shown in panel D. The micrograph shows a CIP2A^wt^ in a dimeric and tetrameric state side-by-side. (**L**) Fraction of events per 2.25 kDa bin of CIP2A^wt^, CIP2A^ΔC^, CIP2A^m4^ particles with determined masses corresponding to the molecular weight of CIP2A dimers and tetramers. plotted as in panel C. For quantification, see Suppl. Fig. 3D.

### CIP2A tetramerizes via an antiparallel coiled coil

Analysis of 401 CIP2A homologs, identified across metazoa (Suppl. Fig. 1), revealed a conserved hydrophobic stretch at the C-terminal end of the predicted parallel coiled coil dimer (Fig. 1F). *In silico* predictions with four copies of the C-terminal region of CIP2A suggest that this stretch, comprising CIP2A^884–892^ in humans, cluster at the core of a short tetrameric antiparallel coiled coil (Fig. 1G). We note that this interaction did not emerge from a query using four copies of full-length CIP2A, highlighting how the visualization of full-length CIP2A via low-resolution electron microscopy inspired a targeted *in silico* structural analysis. To experimentally test the predicted structure, we purified CIP2A^753–905^ and variants that lack the last 5 (CIP2A^ΔC5^), 10 (CIP2A^ΔC10^), or 23 (CIP2A^ΔC23^) residues and compared their multimerization state using analytical size-exclusion chromatography (Fig. 1G). Whereas CIP2A^753–905^, CIP2A^753–900^, and CIP2A^753–895^ formed tetramers, the increased retention volume of CIP2A^753–882^ is consistent with a dimeric state (Fig. 1H, Suppl. Fig 2A). This provides experimental support for the conserved CIP2A^884–892^ [MIAMIHSLS] sequence being at the core of the tetramerization interface. If the prediction of the antiparallel tetrameric coiled coil is accurate, the solvent-exposed S890 should be more permissive to mutation than L891, a residue that contributes to the hydrophobic core of in tetramer state (Fig. 1I). Indeed, mutation of L891 interfered with CIP2A tetramerization (with L891I as a notable exception), whereas mutation of S890 did not (Fig. 1J, Suppl. Fig 2B). Individual mutations within the CIP2A^884–892^ stretch had distinct effects on CIP2A tetramerization, providing further support for the predicted structure (Fig. 1J, Suppl. Fig 2C). To investigate tetramerization-deficient full-length CIP2A, we combined four (M884E, I885N, L891R, S892A; m4) or six (+ M887Q, H889Q; m6) mutations, effectively interfering with the tetramerization of CIP2A^753–905^ (Fig. 1J, Suppl. Fig 3A, B). Tetramerization via the C-terminal motif of CIP2A is further supported by the ability of bead-immobilized human CIP2A^877–905^, but not CIP2A^877–905^ ^m4/m6^, to pull down full-length CIP2A from a *Xenopus* oocyte extract (Suppl. Fig. 3C). Consistently, direct inspection of full-length CIP2A by EM confirmed that tail-tail interactions were abolished in CIP2A^ΔC^ and CIP2A^m4^ mutants (Fig. 1K). A strong reduction of tetramers upon the mutation of the C-terminal region of CIP2A was also observed by mass photometry, and we note that remaining particles with a mass of approximately 400 kDa might be attributed to interdimeric association of the CIP2A globular domains (Fig. 1L, Suppl. Fig 3D)^29^.

### CIP2A tetramerization is required for mitotic DNA repair

We next investigated the impact of CIP2A tetramerization on the recruitment of DNA repair factors onto mitotic chromosomes in RPE1 *TP53*^−/−^ *CIP2A*^−/−^ cells reconstituted with different CIP2A constructs. Following aphidicolin-induced perturbation of DNA replication, we observed that the expression of CIP2A^wt^ resulted in robust co-localizing foci of CIP2A and TOPBP1 on mitotic chromosomes (Fig. 2A-D). In contrast, expression of CIP2A variants with a truncated (CIP2A^ΔC^) or mutated (CIP2A^m4^, CIP2A^m6^) C-terminal tail precluded the formation of distinct CIP2A-TOPBP1 foci at sites of DNA damage (Fig. 2A-D, Suppl. Fig. 4A,B). Likewise, expression of tetramerization-defective CIP2A variants failed to recruit SLX4, a central component of the SMX tri-nuclease complex^23,24^ (Fig. 2E, Suppl. Fig. 4C). Importantly, tetramerization-defective CIP2A still binds TOPBP1 *in vitro* and this requires the previously identified TOPBP1 binding site on the N-terminal globular domain of CIP2A (Suppl. Fig. 4D-F).

**Figure 2:**
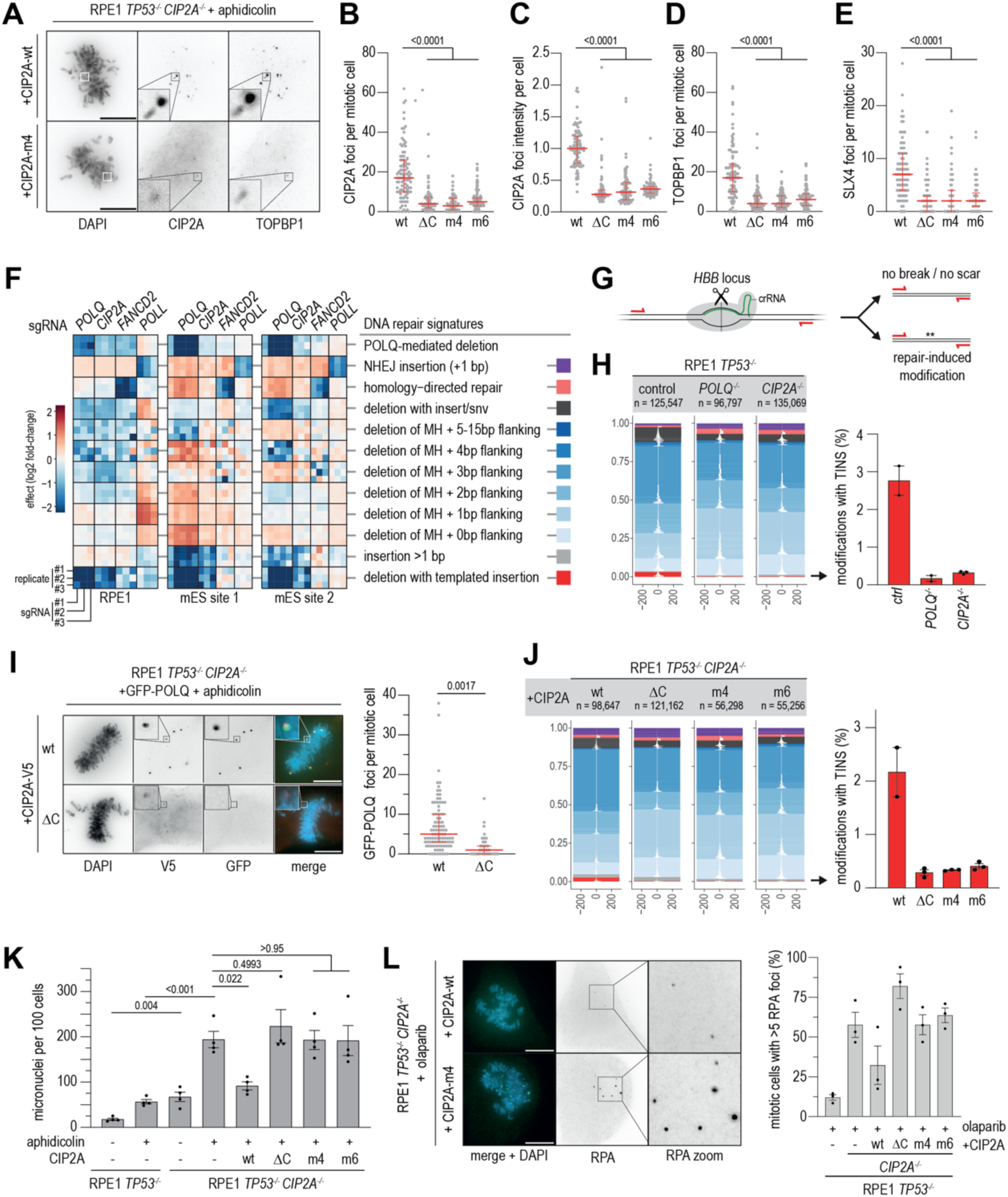
CIP2A tetramerization is required for mitotic DNA repair. **A**) Representative wide-field images of RPE1 *TP53^−/−^ CIP2A^−/−^* cells reconstituted with indicated CIP2A variants and treated with aphidicolin (200 nM). (**B-E**) Quantification of numbers (panel B) or intensity (panel C) of CIP2A foci, or numbers of TOPBP1 foci (panel D) or SLX4 foci (panel E) per mitotic cell. Bars represent medians and interquartile range of three biologically independent experiments, with >90 cells measured per condition across experiments. P-values were calculated via an ordinary one-way ANOVA with Dunnett’s multiple comparisons test on the median per experiment. (**F**) Focused re-analysis of DNA repair signatures in RPE1 or mouse embryonic stem (mES) cells with sgRNAs targeting *POLQ*, *CIP2A*, *FANCD2*, *POLL*. Heatmap represents differences in DNA repair signatures for each target site compared to non-targeting control sgRNAs. Three replicates for at least 3 different sgRNAs per gene are shown. Most relevant repair outcomes are indicated. MH, microhomology. (**G**) Schematic representation of DNA repair outcome analysis upon CRISPR-Cas9 targeting of the *HBB lo*cus. (**H**) Tornado plots of repair outcomes in indicated RPE1 *TP53^−/−^* cells. Percentages of altered reads with deletions with templated insertions (TINS) are indicated. Bars represent means and standard error of the means (SEM) of two (control, *POLQ^−/−^*) or three (*CIP2A^−/−^*) biologically independent experiments. X-axis indicates the distance relatively to the break. (**I**) Wide-field images of RPE1 *TP53^−/−^ CIP2A^−/−^* GFP-POLQ cells reconstituted with indicated CIP2A-V5 variants and treated with aphidicolin (200 nM). Quantification of GFP-POLQ foci per mitotic cell, from 94 cells per condition across three biologically independent experiments. Medians and interquartile range is indicated. A two-tailed unpaired t-test was used. (**J**) Tornado plots of DSB repair products at the *HBB* locus in *CIP2A^−/−^* cells, reconstituted with indicated CIP2A variants. Bars represent means and SEM of two (wt) or three (ΔC, m4, m6) biologically independent experiments. X-axis indicates the distance relatively to the break. (**K**) Quantification of micronuclei per 100 RPE1 *TP53^−/−^ CIP2A^−/−^* cells or RPE1 *TP53^−/−^ CIP2A^−/−^* cells reconstituted with indicated CIP2A-His constructs either left untreated or treated with aphidicolin (200 nM). Bars represent means and SEM of four biologically independent experiments, with >270 cells per condition measured across experiments. P-values were calculated via a two-sided unpaired t-test. (**L**) Representative images of RPE1 *TP53^−/−^ CIP2A^−/−^* cells reconstituted with indicated CIP2A variants, treated with olaparib (0.5 μM) and stained for RPA32. Quantification of the percentage of mitotic cells with >5 RPA32 foci per experiment. Bars represent means and SEM of three biologically independent experiments with >135 cells per condition measured across experiments. Throughout the figure, ‘n’ represents the number of sequencing reads plotted per tornado plot. scalebars represent 10 µm.

To analyse the effects of CIP2A on mitotic DNA repair, we performed a focused analysis of our genome-wide DNA repair signature data, obtained with large sgRNA-libraries in human RPE1 and mouse embryonic stem cells, exploiting genome-covering sgRNA libraries^31^. We note that the inactivation of *CIP2A* or *POLQ* (encoding Polymerase Theta) resulted in remarkably similar DNA repair defects that are distinct from other DNA repair defects, following e.g. the inactivation of *FANCD2* or *POLL* (Fig. 2F). Although the effects of CIP2A depletion on repair outcomes in the category ‘POLQ-mediated deletions’ differed between cell lines, the loss of *CIP2A* or *POLQ* consistently reduced ‘deletions with templated insertions’ (TINS) (Fig. 2F). Since TINS are an exclusive hallmark of DNA repair via POLQ (Polymerase Theta)-mediated end joining (TMEJ)^32^, this provides direct evidence for CIP2A-mediated TMEJ during mitosis.

To investigate in more detail how CIP2A impacts DNA repair outcomes, we used CRISPR/Cas9 to induce DSBs in the *HBB* locus in RPE1 *TP53^−/−^*cells with or without *CIP2A* and *POLQ* (Fig. 2G). We found that in addition to inactivation *POLQ,* also genetic inactivation of *CIP2A* resulted in TMEJ defects, characterized by fewer deletions flanked by microhomology and fewer deletions with insertions (Suppl. Fig. 5A), as well as drastically fewer TINS (Fig. 2H). Consistently, and in line with previous studies^24,28^, GFP-POLQ formed discernible mitotic foci in control cells, but not in *CIP2A^−/−^* cells (Suppl. Fig. 5B,C). To test how CIP2A tetramerization impacts mitotic DNA damage processing, we studied the localization of POLQ in *CIP2A*^−/−^ cells that expressed tetramerization-defective CIP2A^ΔC^. Consistent with the key role of CIP2A self-interaction via the C-terminal tail, we find a significant reduction in GFP-POLQ foci at mitotic DNA lesions, demonstrating that the C-terminal region of CIP2A is required for the efficient recruitment of POLQ (Fig. 2I, Suppl. Fig. 5D). Consequently, the expression of tetramerization-defective CIP2A variants resulted in defective DNA repair. Specifically, while TINS were abolished in *CIP2A^−/−^*cells (Fig. 2H), these repair products re-appeared upon expression of CIP2A^wt^, but not upon the expression of CIP2A with a truncated (CIP2A^ΔC^) or mutated C-terminal tail (CIP2A^m4^, CIP2A^m6^) (Fig. 2J). The expression of tetramerization-defective CIP2A variants also failed to rescue other TMEJ repair products, including deletions with microhomology and deletions with insertions (Suppl. Fig. 5E). Combined, our data show that CIP2A tetramerization is required for mitotic DNA repair by POLQ.

### CIP2A prevents RPA accumulation and micronucleation

A failure to properly process and tether mitotic DNA lesions results in the formation of micronuclei^17,16,23,30,33^. Indeed, *CIP2A* inactivation leads to elevated levels of micronuclei in unperturbed conditions, which are further elevated upon aphidicolin-induced DNA damage (Fig. 2K). Notably, whereas expression of CIP2A^wt^ suppressed the aphidicolin-induced micronucleation in *CIP2A^−/−^* cells, expression of tetramerization-defective CIP2A variants (CIP2A^ΔC^, CIP2A^m4^ or CIP2A^m6^) failed to reduce micronucleation (Fig. 2K).

Mitotic DNA breaks, either directly induced during mitosis or resulting from processing of incompletely replicated DNA, only represent a subset of mitotic DNA lesions. To assess whether CIP2A tetramerization is required for the response to other forms of replication-mediated mitotic DNA damage, we investigated RPA foci formation. The number of RPA foci, marking single-stranded DNA, increased significantly in *CIP2A*^−/−^ cells (Suppl. Fig 5F). Upon treatment with olaparib, a PARP inhibitor that is known to cause persistent single stranded DNA gaps^5^, the number of RPA foci in *CIP2A*^−/−^ cells increased further (Suppl. Fig 5G). This increase was reduced upon the expression of CIP2A^wt^, but not upon the expression of tetramerization-defective CIP2A variants (Fig. 2L, Suppl. Fig 5G). Combined, these data show that CIP2A tetramerization is required for the recruitment of DNA repair factors at mitotic DNA lesions and for their repair.

Of note, the C-terminal tail of CIP2A harbors a previously reported phosphorylation site (S904) that matches a PLK1 consensus motif and affects spindle pole localization and recruitment of TOPBP1 and MDC1 in oocytes^34,35^. Since both CIP2A (Suppl. Fig. 5C) and PLK1 activity are required for POLQ recruitment to mitotic DNA lesions^24,28^, we explored the relevance of this putative regulatory site in detail. However, non-phosphorylatable and phosphorylation-mimicking mutants (CIP2A^S904A/D/E^) were both indistinguishable from CIP2A^wt^ regarding TOPBP1 and GFP-POLQ recruitment, DNA repair signatures, and micronuclei formation (Suppl. Fig. 6A-G). Consistently, CIP2A^S904A^ and CIP2A^S904D^ also showed tetramer formation comparable to CIP2A^wt^ *in vitro* (Suppl. Fig. 6H-J).

### CIP2A tetramerization is a conserved and transferable trait

To investigate the structural features of CIP2A across metazoans, we analysed coiled coil patterns and structural predictions for CIP2A homologs *in silico*. This comparison showed that the dimerization via near-continuous long parallel coiled coils is a conserved feature of CIP2A (Fig. 3A) and that, despite a striking variation at the sequence level, four copies of CIP2A are consistently predicted to tetramerize via a short antiparallel coiled coil (Fig. 3B,C, Suppl. Fig 7A). After establishing that tetramerization-deficient forms of CIP2A do not support foci formation of DNA repair factors during mitosis (Fig. 2), we hypothesized that other tetramerization interfaces at the C-terminal end of CIP2A might suffice to restore mitotic DNA repair. To test this, we replaced human CIP2A^870–905^ with C-terminal tails from CIP2A homologs found in *A. cephalotes*, *C. elegans*, *D. melanogaster*, and *O. bimaculoides*, carefully maintaining the coiled coil register (Fig. 3C, Suppl. Fig 7B). In an orthogonal approach, inspired by the architecture of tetrameric coiled coils^36^, we introduced three heptads of a synthetic antiparallel tetrameric coiled coil at the C-terminal end of CIP2A (Fig. 3D). Size exclusion chromatography of all five CIP2A chimeric fragments demonstrated their ability to tetramerize *in vitro* (Fig. 3E, Suppl. Fig 7C, D). Importantly, tetramerization of the CIP2A chimera fragments critically relied on Leucine residues, equivalents of the human CIP2A^L891^, positioned at the core of the tetramerization domain (Fig. 3E, Suppl. Fig 7C). This analysis provides further experimental support for the predicted structural models and highlights the remarkable divergence at the sequence level in CIP2A tetramerization motifs across metazoans.

**Figure 3:**
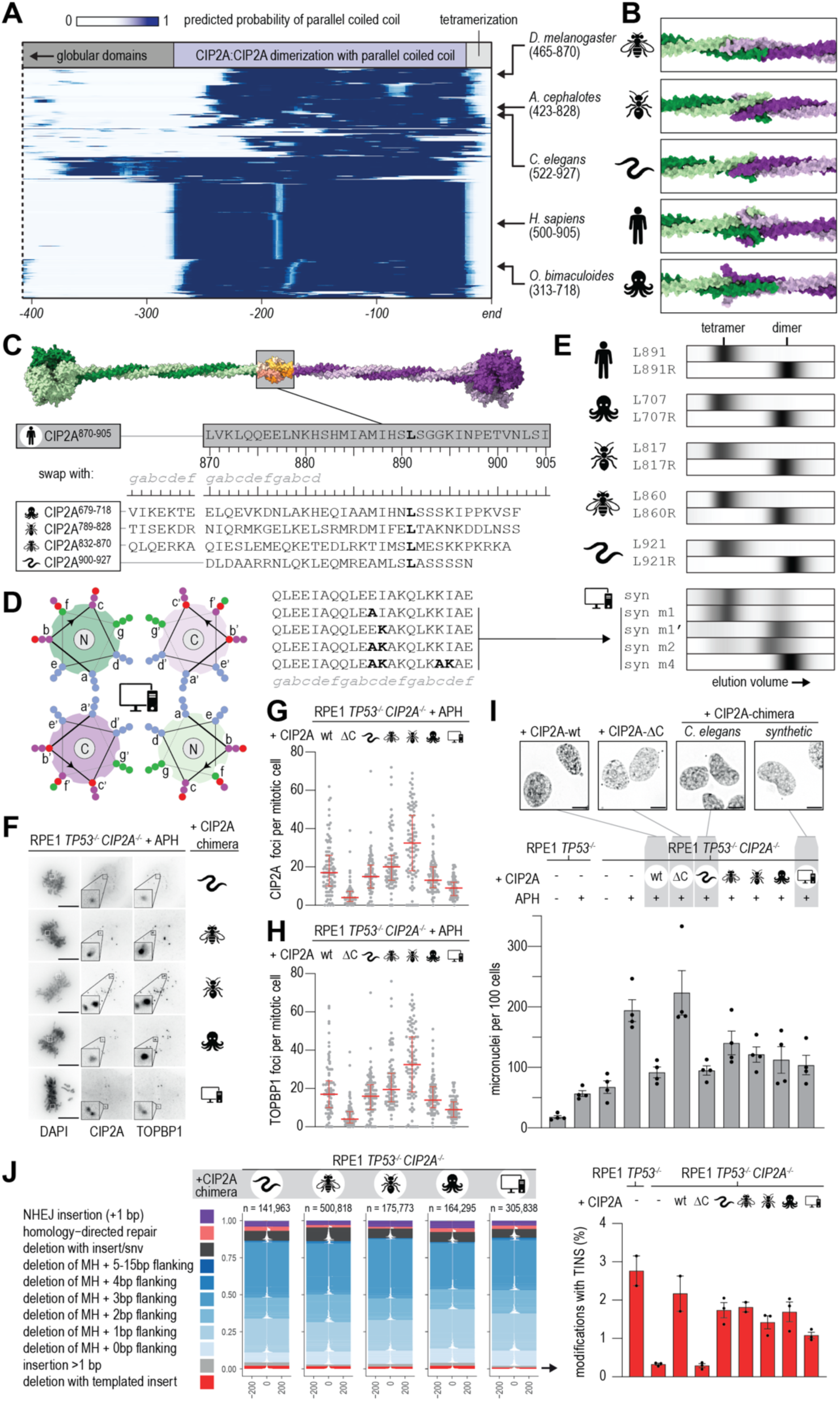
CIP2A tetramerization is a conserved and transferable trait. (**A**) CIP2A homologs (n = 434) were scored for their probability to form a parallel coiled coil (CC). Their most C-terminal 405 residues are shown here, with dark blue representing a high-confidence CC, after they were clustered hierarchically based on the per-residue CC probability scores. Arrows indicate the tracks of the selected species and their last 405 residues. (**B**) Structural predictions of the C-terminal regions of four copies in the selected species. (**C**) Structural prediction-based cartoon of four human CIP2A^1–905^ copies, as in Figure 1G. Grey boxes show the region and the sequence of the human CIP2A^870–905^ stretch that is replaced with the displayed sequences from CIP2A homologs. Equivalents of the human Leucine 891 are shown in bold. (**D**) Helical wheel representation of a tetrameric antiparallel CC, as in Fig. 1I, with three heptads of a synthetic sequence. Bold residues in the displayed sequences indicate mutations that are predicted to interfere with tetramerization. (**E**) Size-exclusion chromatography analysis of CIP2A^753–905^ constructs with all indicated chimeric variants and mutations, summarized as in Figure 1H. (**F**) Representative wide-field images of RPE1 *TP53^−/−^ CIP2A^−/−^* cells reconstituted with indicated CIP2A variants stained for DAPI, CIP2A and TOPBP1 after aphidicolin treatment (200 nM).(**G**-**H**) Quantification of the number of CIP2A foci and TOPBP1 foci per mitotic cell. Bar represents the median and interquartile range of three biological independent experiments with at least a total of 90 cells measured per condition across experiments. Wt and ΔC are identical to the samples plotted in Fig. −28C. (**I**) Representative images of RPE1 *TP53^−/−^ CIP2A^−/−^* cells reconstituted with indicated CIP2A variants stained for DAPI after aphidicolin treatment (200 nM). Quantification of the number of micronuclei per cell in RPE1 *TP53^−/−^ CIP2A^−/−^* cells and RPE1 *TP53^−/−^ CIP2A^−/−^* cells reconstituted with indicated CIP2A variants either left untreated or treated with aphidicolin (200 nM). Bars represent the mean and SEM of four biologically independent experiments with at least a total number of 265 cells per condition measured across experiments. RPE1 *TP53^−/−^ CIP2A^−/−^* cells with or without CIP2A wt or ΔC are identical to the samples plotted in Fig. 2K. (**J**) Tornado plots of repair outcomes for RPE1 *TP53^−/−^ CIP2A^−/−^* cells reconstituted with indicated CIP2A variants. Percentages of altered reads with deletions with TINS are indicated. Bars represent the mean and SEM of two (RPE1 *TP53^−/−^*, wt and *D. melanogaster*) or three (*CIP2A^−/−^*, ΔC, *C. elegans*, *A. cephalotes*, *O. bimaculoides* and synthetic) biologically independent experiments. RPE1 *TP53^−/−^ CIP2A^−/−^*, with or without CIP2A wt or ΔC, are idenntical to the samples plotted in Fig. 2. ‘n’ represents the number of reads plotted per tornado plot. X-axis indicates the distance relatively to the break. Throughout the figure, scalebars represent 10 µm.

To test if CIP2A chimeras are functional in cells, we reconstituted RPE1 *TP53*^−/−^*CIP2A^−/−^* cells with the chimeric CIP2A variants described above (Suppl. Fig. 4A) and tested their ability to support the recruitment of CIP2A, TOPBP1 and SLX4 to mitotic DNA lesions. Importantly, all CIP2A chimeras formed CIP2A foci upon aphidicolin exposure (Fig. 3F, G). These foci also contained TOPBP1 (Fig. 3F, H), and SLX4 (Suppl. Fig. 8A). Although we observed variation among CIP2A chimeric constructs, as reflected in varying levels of γH2A.X foci (Suppl. Fig. 8B), a comparison with cells expressing CIP2A^ΔC^ demonstrated that all sequences restored the functionality of CIP2A at least partially (Fig. 3F-H) and were able to suppress aphidicolin-induced micronuclei (Fig. 3I). To explore if CIP2A chimeras also facilitated mitotic DNA repair, we analysed DNA repair profiles at CRISPR/Cas9-mediated breaks at the *HBB* locus. Consistent with successful restoration of CIP2A and TOPBP1 foci, all CIP2A chimeras restored repair outcomes, particularly those involving TINS (Fig. 3J, Suppl. Fig. 8C). These data demonstrate that the tetramerization of CIP2A via a short antiparallel coiled coil is a functionally conserved and transferrable element that supports mitotic DNA repair.

### CIP2A tetramerization promotes fitness of BRC-1 deficient *C. elegans*

To investigate the importance of CIP2A for DNA repair during development, we examined mutants in ORF B0432.7 in *C. elegans*, which encodes a *bona fide* but uncharacterized CIP2A homolog. We generated a strain lacking the entire ORFB0432.7, hereafter called *cip2-A^ko^* and a strain with tetramerization-defective CIP-2A, lacking amino acids 914-922, hereafter called *cip-2A*^Δ*C*^ (Suppl. Fig. 9A,B). Both *cip2-A^ko^* and *cip-2A*^Δ*C*^ did not affect brood sizes and hatching rates of the embryos (Fig 4A,B). This indicates that neither mitotic nor meiotic problems occurred in the absence of CIP-2A. Inspired by the finding that TMEJ via POLQ promotes the survival of *C. elegans* in the absence of BRC-1^37^, the orthologue of human BRCA1, we set out to investigate the effects of *cip-2A^ko^*and *cip-2A*^Δ*C*^ in a *brc-1* null background. This revealed mild but reproducible synthetic effects of *cip-2A* and *brc-1* on brood size and offspring viability (Fig. 4A, B). The comparable and significant decreases in brood size and embryonic viability in the *cip-2A*^Δ*C*^ and *cip2-A^ko^*alleles in combination with *brc-1* corroborates the importance of CIP2A’s C-terminal region.

**Figure 4:**
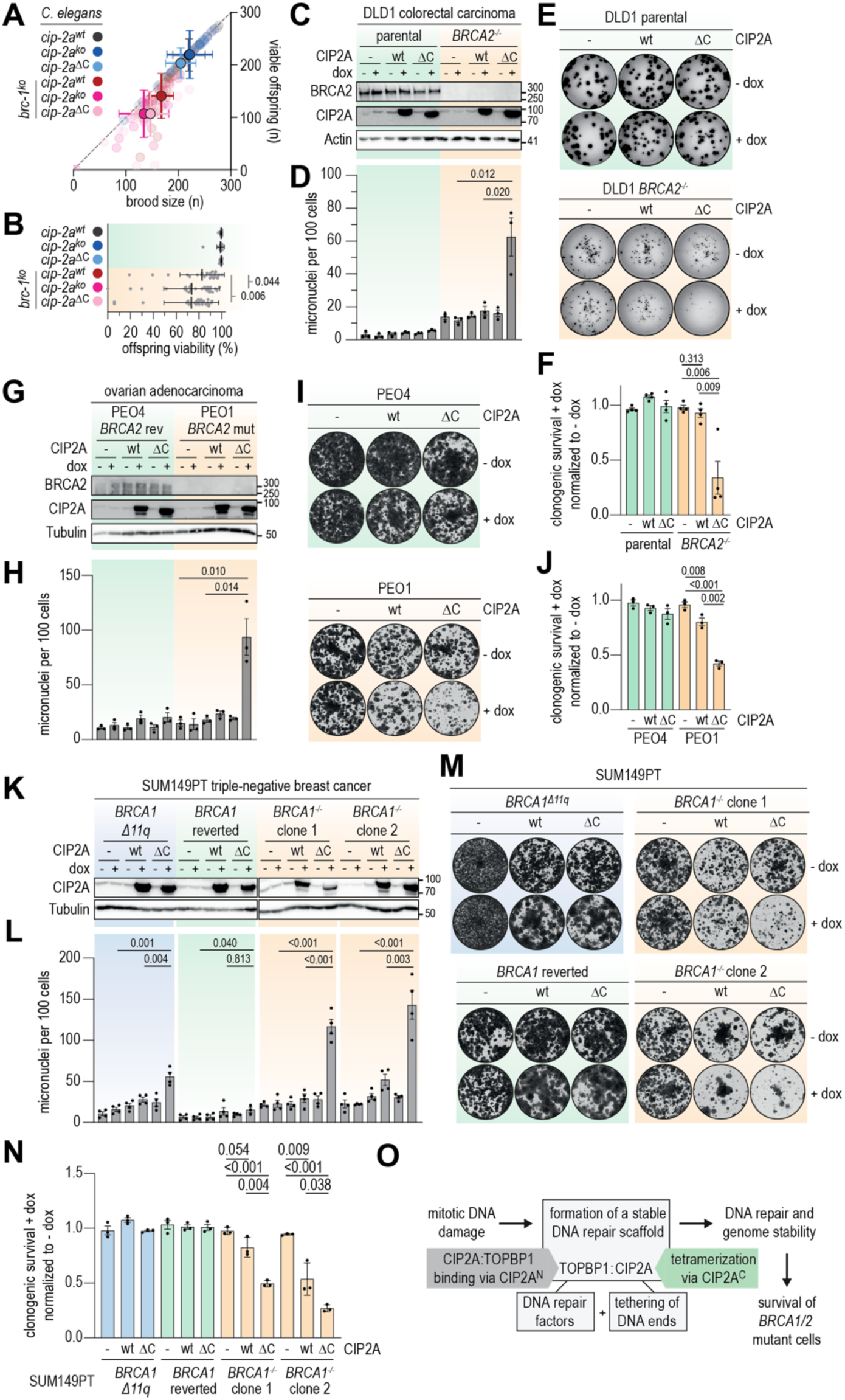
Expression of CIP2A DC exerts dominant-negative effects and impairs survival of *BRCA1/2* mutant cells. **(A)** The number of laid and hatched eggs was followed over 3 days from 25 hermaphrodites for the indicated genotypes. Brood size is the number of laid eggs per hermaphrodite mother animal; viable offspring indicates the number of hatched eggs per hermaphrodite. Bars represent the mean and standard deviation SD). (**B**) Percentage of hatched eggs per hermaphrodite per genotype. Bars represent the mean and SD. P-values were calculated via a two-sided Mann-Whitney U test. (**C, D**) Parental and *BRCA2^−/−^* DLD1 cells, stably expressing doxycycline-inducible CIP2A^wt^ or CIP2A^ΔC^ were treated with doxycycline (dox, 0.1 mg/mL) for 48 hours. Immunoblot analysis of indicated proteins and micronuclei per 100 cells are indicated. Bars represent means and SEM of at least three biologically independent experiments, with in total >956 cells per condition. **(E, F)** Representative images and quantification of clonogenic survival of indicated DLD1 cell lines. Bars represent the mean and SEM of four independent experiments. **(G, H)** PEO4 and PEO1 cells stably expressing doxycycline-inducible CIP2A^wt^ or CIP2A^ΔC^ were treated with doxycycline (1 mg/mL) for 48 hours. Immunoblot analysis of indicated proteins and micronuclei per 100 cells are indicated. Bars represent means and SEM of at least three biologically independent experiments, with in total >437 cells per condition. **(I, J)** Clonogenic survival of PEO4/1 cell lines. Bars represent the mean and SEM of three independent experiments. **(K, L)** Immunoblotting and micronuclei analysis of parental BRCA1 D11q), *BRCA1* reverted, *BRCA1^−/−^* cl#1 and *BRCA1^−/−^* cl#2 SUM194PT cells, expressing CIP2A^wt^ or CIP2A^ΔC^. **(M, N)** Clonogenic survival analysis of indicated SUM149PT cells. Bars represent the mean and SEM of at least three biologically independent experiments with in total >620 cells per condition. Two-tailed unpaired t-tests were used for panels D-N. **(O)** Schematic representation of the role of CIP2A tetramerization in mitotic DNA repair scaffold formation and mitotic DNA repair and survival of *BRCA1/2* mutant cells.

### Tetramerization-defective CIP2A is dominant negative

Human cells with mutations in *BRCA1* or *BRCA2* strictly rely on CIP2A for their survival^17,23,16^. To investigate if tetramerization-deficient forms of CIP2A selectively sensitize human cells with homologous recombination (HR) deficiencies, we expressed CIP2A mutants in the presence of endogenous CIP2A (Suppl. Fig. 10A). The expression of CIP2A^ΔC^ in RPE1 *TP53*^−/−^ cells resulted in fewer and less intense CIP2A, TOPBP1, SLX4, and POLQ foci in mitotic cells (Suppl. Fig. 10A-H). This was not observed upon the expression of CIP2A^wt^ (Suppl. Fig. 10A-H), indicating that CIP2A^ΔC^ impaired the mitotic DNA damage response in a dominant-negative fashion. Consistently, high levels of CIP2A^ΔC^ promoted the formation of micronuclei, especially in the presence of aphidicolin-induced DNA replication stress (Suppl. Fig. 10I). Based on these findings, we hypothesized that the presence of CIP2A^ΔC^ could selectively decrease the viability of cells with high levels of mitotic DNA damage. To test this, we expressed CIP2A^wt^ or CIP2A^ΔC^ in DLD1 *BRCA*2^−/−^ cells (Fig 4C), previously demonstrated to have high levels of mitotic DNA lesions^17,23,38^. After 4 hours of doxycycline-induced CIP2A^ΔC^ expression in DLD1 *BRCA2^−/−^* cells, mitotic CIP2A and TOPBP1 foci were reduced (Suppl. Fig. 11A,B). Prolonged expression (48 h) further decreased CIP2A, TOPBP1, and SLX4 mitotic foci (Suppl. Fig. 11C-F). Moreover, expression of CIP2A^ΔC^ significantly increased the number of micronuclei in DLD1 *BRCA2^−/−^* cells (Fig. 4D, Suppl. Fig. 11G) and reduced clonogenic survival (Fig. 4E).

Importantly, neither overexpressing CIP2A^ΔC^ in *BRCA2^+/+^*cells nor overexpressing CIP2A^wt^ in *BRCA2^+/+^* or *BRCA2^−/−^* DLD1 cells influenced clonogenic survival (Fig. 4E, F). To extend these findings to other HR-deficient models, we overexpressed CIP2A^wt^ or CIP2A^ΔC^ in ovarian adenocarcinoma cells with a *BRCA2* 4965C>G mutation (PEO1) and a derivative isogenic HR-proficient model with a *BRCA2* 4965C>T reversion mutation (PEO4)^39^ (Fig. 4G). We also investigated the effect of high CIP2A^ΔC^ levels in SUM149PT triple-negative breast cancer cells harboring a *BRCA1* mutation (2288delT), SUM149PT cells with a *BRCA1* reversion mutation, and two SUM149PT clones with a CRISPR/Cas9-mediated complete loss of *BRCA1* (Fig. 4K, Suppl. Fig. 11H). CIP2A^ΔC^ expression resulted in significantly increased numbers of micronuclei and reduced clonogenic survival in the *BRCA2*-deficient PEO1 cells (Fig. 4G-J, Suppl. Fig. 11I) and the SUM149PT *BRCA1^−/−^*cells (Fig. 4K-N, Suppl. Fig. 11J). Taken together, our results demonstrate that overexpression of the tetramerization-defective CIP2A^ΔC^ variant disrupts the function of endogenous CIP2A, preventing the formation of mitotic DNA repair foci and leading to increased genome instability and loss of viability in BRCA1/2 deficient cells.

## Discussion

Here, we demonstrate that CIP2A forms elongated tetramers that contribute to genome stability by promoting POLQ-mediated DNA repair during mitosis (mitotic TMEJ). HR-deficient cells rely on this pathway for their survival^40,41,27,28^, explaining their hypersensitivity to CIP2A perturbations^16,17,23^. We propose that the tetramerization of CIP2A is required to form a stable DNA repair scaffold that effectively coordinates the recruitment of DNA processing factors and the tethering of DNA ends (Fig. 4O). How tetramerization coordinates these activities remains incompletely clear from a molecular perspective. It is known that CIP2A exerts its effect on DNA repair via TOPBP1 and that this requires the binding of the N-terminal domain of CIP2A to an unstructured TOPBP1 region positioned between TOPBP1’s BRCT5 and BRCT6 domains^17,42^ (Suppl. Fig. 4F). Importantly, upstream factors that mediate TOPBP1-CIP2A binding to replication-mediated DNA lesions remain elusive. Possible candidates include the Rad9-Rad1-Hus1 (9-1-1) complex^43–45^, in conjunction with RAD17^46^ and RHNO1^47,27^, which were previously identified to interact with TOPBP1 and to be recruited to stalled replication forks. Alternatively, binding of TOPBP1 to RPA at single-stranded DNA may be involved in TOPBP1-CIP2A recruitment to mitotic DNA lesions^48^.

With TOPBP1-DNA interactions at either end of a dumbbell-like CIP2A tetramer, our findings provide a molecular glimpse of a mitotic DNA repair scaffold and have a number of implications for the putative contribution of CIP2A to mitotic DNA tethering and repair. First, CIP2A multimers might efficiently and locally increase the concentration of TOPBP1 and its downstream factors. Second, the elongated nature of CIP2A could serve as a molecular spacer and impose geometric constraints, positioning DNA lesions relative to effector proteins, to promote productive DNA repair. Third, CIP2A multimers might provide an extra layer of avidity to a DNA repair scaffold, possibly important in the context of condensed mitotic chromosomes. A similar mechanism might be at play following the shattering of micronuclei when CIP2A has been shown to tether damaged DNA ends in the cytoplasm during interphase^21,22^.

The near-continuous coiled coils of CIP2A and the tail-tail tetramerization results in particles of almost 100 nm. Interestingly, dumbbell-like CIP2A tetramers with TOPBP1-mediated DNA processing activities at their ends resemble the appearance of other DNA tethering protein complexes, including the Mre11-Rad50-Nbs1 complex, the bacterial SbcCD complex^49–51^, CtIP^52,53^ and the synaptonemal complex (SC)^54–56^. In this context, it is noteworthy that we observed occasional interactions between the N-terminal globular domains of CIP2A dimers *in vitro* (Fig. 1L, Suppl. Fig. 3D), possibly providing a molecular basis for further self-interactions and for the previously observed higher-order complexes of TOPBP1-CIP2A in cells^17,23^.

The DNA repair defects in CIP2A^−/−^ cells or in cells expressing tetramerization-defective CIP2A variants involved a near-complete loss of deletions with templated insertions, resembling signatures in *POLQ*^−/−^ cells. Of note, other POLQ-mediated DNA repair signatures were less affected by the absence of CIP2A (*i.e*. deletions with microhomology), then by the absence of POLQ (Suppl. Fig. 5D), supporting a model in which POLQ functions in CIP2A-dependent and CIP2A-independent DNA repair. These different repair outcomes likely reflect mitotic and interphase roles of POLQ, respectively. Although chimeric CIP2A variants differed slightly regarding DNA repair outcomes (Suppl. Fig. 8C), they effectively rescued micronucleation (Fig. 3I). This demonstrates that tetramerization of CIP2A C-termini *per se* is essential to guide DNA repair during mitosis.

It is likely that TOPBP1-CIP2A is also required for DNA repair of lesions that do not involve DSB formation. Consistently, only a subset of mitotic DNA lesions recruit POLQ and the SMX complex^23^. Furthermore, persistent ssDNA gaps were recently also demonstrated to be transmitted into mitosis and require TOPBP1-CIP2A for processing^57,58^. In line with this notion, we observed that cells lacking CIP2A or expressing tetramerization-defective CIP2A variant show persistent RPA foci in response to PARP inhibition, pointing at a general requirement for CIP2A tetramerization in mitotic DNA repair.

CIP2A was previously identified as a synthetic lethal target in *BRCA1/2* mutant cancer cells^17^. However, multiple genotoxic drugs induce DNA lesions that are transmitted into mitosis ^59^, indicating a broader dependency on CIP2A in cancer cells. Our data show that tetramerization-defective CIP2A mutant exert dominant-negative effects, and their expression consequently impairs the clonogenic survival of HR-defective cancer models. Yet, reversion mutations that are known to restore HR nullified sensitivity to disrupted CIP2A tetramerization. This highlights HR-deficiencies as the key determinant for CIP2A dependency and warrants studies that explore inhibition of CIP2A tetramerization as a possible therapeutic strategy.

## Author contributions

LdH, MATMvV, and PJH conceived the study. RS, EO, SSB, and PJH performed biochemical assays. PJH performed structure-function analysis and electron microscopy. LdH, MLK, FJB, and HRB performed cell biological experiments, supervised by JJ and MATMvV. DNA repair assays were performed by LdH, FJB, HRB, MB, and AD, supervised by MT. AG and DP performed *C. elegans* experiments, supervised by VJ. BvdK and RF developed reagents. LdH, MATMvV and PJH wrote the manuscript and all authors provided feedback.

## Acknowledgements

This work was financially supported by the Dutch Cancer Society (#12911 to MATMvV), the Netherlands Organization for Scientific Research (NWO VICI #09150182110019 to MATMvV), and the Austrian Science Fund (FWF PAT1893225, https://doi.org/10.55776/PAT1893225, to PJH and SFB F8805-B, https://doi.org/10.55776/F88, to VJ). Electron microscopy was performed at the Electron Microscopy Facility of the Vienna BioCenter Core Facilities (VBCF), member of the Vienna BioCenter (VBC), Austria. We acknowledge Nicola Silva for the *brc-1(ddr41) C. elegans* strain. We are grateful to Nellie Prinz, Marie Stute, and Iulia Nicolaescu for help with cloning and protein purification, Puck Knipscheer for mitotic *Xenopus laevis* egg extracts, Alexander Schleiffer for discussions, Robin van Schendel and Joost Schimmel for help with DNA signature analysis, and Marieke Everts and Marlene Brandstetter for technical support. We thank Joao Matos and Gerben Vader for discussions and commenting on the manuscript.

## Methods

### Sequence alignments, structural predictions, and coiled-coil analysis

Multiple sequence alignments were generated from 490 CIP2A sequences from Ophistokonta, collected from Eggnog6^59^ (7MGG7) complemented with 86 sequences from oomycota, curated on InterPro^60^ (PF21044). Duplicates were removed and sequences were manually filtered using Jalview^61^ for sequence quality and completeness after an alignment via ClustalW^62^. Structures of CIP2A homologs were inspected from in silico prediction databases (for monomers), predicted for selected species using Alphafold3^63^ (two full-length copies, four copies of the most C-terminal 100-150 residues), and analysed using ChimeraX^64^ and PDBePISA^65^.

From the remaining 442 sequences, the last 700 residues were analysed for their predicted likelihood to form a parallel dimeric coiled coil via CoCoNat^66^. After filtering out 8 sequences of unidentified species or without recognizable CIP2A coiled coil (CC) patterns, 434 sequences remained. Predicted CC registers were used to accurately replace the C-terminus of human CIP2A with sequences from homologs or synthetic sequences. To make the CC diagrams, these 434 CIP2A homologs were scored for their propensity to form a parallel dimeric CC and coloured per-residue. Sequences were aligned on their C-termini to allow comparisons across sequences of varying total length. Sequences shorter than 700 residues were zero-padded on the left. The CC probability profiles in the last 350 C-terminal residues were used to cluster sequences hierarchically according to Ward’s minimum variance method using Euclidean distance as the distance metric via the scipy-based seaborn.clustermap implementation. Sequences were classified taxonomically using the NCBI taxonomy API and the UniProt REST API.

For C-terminal alignments, the 401 sequences with a maximum 50 residues after the last heptad of the parallel dimeric CC were used. Starting 70 residues prior to this last heptad, sequences were aligned using MAFFT^67^ with a high hap penalty. Alignments were inspected in Jalview. From 401 sequences, a sequence logos was generated using Logomaker^68^, with letter heights reflecting Shannon information content calculated from per-column amino acid frequency data exported from Jalview. Letter transparency was scaled by per-column occupancy. Alignment colours follow the ClustalX amino acid colour scheme throughout. Helical wheel representations of antiparallel tetrameric coiled coils were drawn in adobe illustrator based on predicted structures^65,64,63^ with inspiration from https://grigoryanlab.org/drawcoil/^69^ and coiled coil analyses^36,70^.

### Expression and purification of full-length CIP2A

Full-length human CIP2A was cloned into a pLIB vector with a combined sortase-polyhistidine tag (GLPETGGGHHHHHH) at its C-terminal. Mutations and truncations were introduced using *in vivo* assembly (IVA)^71^ and validated via Sanger sequencing. Constructs were transposed into DH10^Embac-YFP^ cells^72^ and extracted bacmids were delivered into *Sf9* insect cells via FuGene HD Transfection Reagent (Promega). All insect cells were cultured at 27°C in ESF 921 (Expression systems). Transfected cells were grown in 6-well plates for 5 days before virus-containing supernatant was used to infect *Sf9* cells in suspension cultures of 25 mL. After 5 days of culturing, the cells were spun down at 3000 rpm for 10 minutes and the supernatant was filtered through 0.2 µm membranes. 1 mL of this virus-containing supernatant was used to infect 25 mL pre-cultures of *Sf9* cells for 3.5 days before 10 mL of pre-culture was added to a 250 mL expression culture of High Five cells. Infected cells were cultured for 2.5 days and harvested by centrifuging at 3000 rpm for 10 min, washing with PBS, and flash-freezing the cell pellet in liquid nitrogen for storage at −70°C.

For Ni-NTA-based affinity chromatography, all steps were performed on ice or at 4°C. Insect cell pellets from a 250 mL culture were resuspended in 20 mL lysis buffer (500 mM NaCl, 50 mM Tris-HCl pH 6.8, 10% (v/v) Glycerol, 15 mM BME, 10 mM Imidazole, 0.1% (v/v) Tween-20, 1 mM PMSF) supplemented with protease inhibitor cocktail (cOmplete, Roche). Cells were lysed by sonication, and the lysate was centrifuged at 20 000 g for 45 min. The soluble fraction was incubated with 1 mL of equilibrated Ni-NTA agarose beads (Protino, Machery-Nagel) for at least 1 hour. Ni-NTA beads were washed twice with buffer (500 mM NaCl, 50 mM Tris-HCl pH 6.8, 5% (v/v) Glycerol, 15 mM BME, 40 mM Imidazole), followed by elution with buffer containing 400 mM Imidazole. For washes, beads were centrifuged at 500 *g* for 5 min. Eluates were loaded on top of a 4 mL glycerol gradient (10-35% v/v glycerol in buffer containing 50 mM Tris pH 6.8, 500 mM NaCl and 2 mM TCEP), prepared using a Gradient Master (BioComp Instruments). The glycerol gradients were ultra-centrifuged in a pre-cooled Sw60Ti rotor (Beckmann) at 200 000 g overnight. Fractions of 350 µL were collected manually from top to bottom and analysed by SDS-PAGE and Coomassie staining. Fractions 4 and 5 were selected for subsequent experiments, i.e. electron microscopy and mass photometry. Fractions were snap frozen in liquid nitrogen and stored at −70°C.

### Expression and purification of the CIP2A coiled coil

The second half of the CIP2A coiled coil, spanning residues 753-905, was inserted into a pOPIN-B derived vector (kindly provided by the Plaschka lab, IMP Vienna) to generate a construct with an N-terminal GST-PreScission tag and a C-terminal sortase-polyhistidine tag (GLPETGGGHHHHHH). Mutations and truncations were introduced using *in vivo* assembly (IVA)^71^ and validated via Sanger sequencing. Constructs were transformed into *E. coli* BL21-CodonPlus(DE3)-RIL cells for protein expression. Cultures were inoculated from a single colony and grown at 37°C in LB in the presence of Kanamycin and Chloramphenicol. After induction at an OD_600_ of 0.6 with 300 µM IPTG, cells were kept at 16°C overnight, harvested by centrifugation (3.000 *g*, 10 minutes, 4°C), and washed with cold PBS. Cell pellets were flash-frozen in liquid nitrogen and stored at −70°C. All subsequent steps were performed on ice or at 4°C. Cell pellets were resuspended in 10 mL lysis buffer (500 mM NaCl, 50 mM Tris-HCl pH 7.4, 10% (v/v) Glycerol, 20 mM BME, 10 mM Imidazole, 0.1% (v/v) Tween-20, 1 mM PMSF) supplemented with complete protease inhibitor cocktail (Roche). Cells were lysed by sonication and centrifuged at 20.000 *g*. Soluble material was incubated with approximately 100 µL equilibrated glutathione agarose beads (Protino, Machery-Nagel) per 100 mL of bacterial culture for at least 3 hours. Glutathione beads were pelleted by centrifugation at 600 *g* for 5 min. After two washes with 10 mL of wash buffer (500 mM NaCl, 50 mM Tris-HCl pH 7.4, 5% glycerol (v/v), 5 mM BME), beads were resuspended in a 1:1 bead:buffer ratio and incubated over-night in the presence of GST-PreScission-3C protease (Huis lab). After pelleting the beads by centrifugation, the supernatant containing CIP2A was loaded onto columns for preparative or analytical size exclusion chromatography (SEC). Preparative SEC was performed on an ÄKTA pure system (Cytiva) with a Superdex 200 Increase 10/300 column (Cytiva) equilibrated in SEC buffer containing 20 mM HEPES pH 7.4, 2% glycerol (v/v), 400 mM NaCl, 1 mM TCEP. Protein elution was followed using absorbance at 215 nm. Selected fractions were analysed by SDS-PAGE and Coomassie staining, snap frozen in liquid nitrogen, and stored at −70°C.

### Analytical size exclusion chromatography

For analytical size exclusion chromatography Short coiled-coil CIP2A constructs, containing residues 753-905 and variants thereof, were purified by affinity chromatography and applied to Superdex 200 Increase columns (Cytiva) with dimensions 10/300 (Suppl. Figure 3A) or 5/150 (other data). Analytical SEC was performed on an ÄKTA PURE micro system (Cytiva). Proteins were analysed by SDS-PAGE and Coomassie staining to ensure comparable protein purity and amounts and injected at CIP2A concentrations of 5-50 µM (calculated as a monomer). Columns were equilibrated in 20 mM HEPES pH 7.4, 2% glycerol (v/v), 400 mM NaCl, 1 mM TCEP. Protein elution was followed using absorbance at 215 nm. Elution profiles were visualized as greyscale strips using a python-based script. In brief, absorbance values were first normalized according to the maximum value within the depicted range of elution volume, next used to calculate the area under the curve and subsequently adjusted corrected according to the sample within a set of measurements with the lowest area under the curve. Fractions of 750 µL (10/300) or 80 µL (5/150) were collected and analysed using SDS-PAGE and Coomassie staining.

### Mass photometry

Mass photometry was performed on a Refeyn Two^MP^ system. Calibration was done with BSA (66.5 and 133 kDa) and Thyroglobulin (330, 660, and 1320 kDa). Glycerol gradient fractions were pre-diluted with PBS to a concentration of approximately 0.5 µM and kept on ice. Final dilutions in PBS to 50 nM were made directly on the slide in the wells of the gasket. Each sample was measured directly after diluting and for the duration of 1 minute. For each measurement a new pre-dilution was made, and a fresh well was used. Data was exported from the Discover^MP^ software (Refeyn) and plotted and analysed using Python-based scripts.

### Metal shadowing and electron microscopy

Freshly harvested CIP2A-containing fractions from a glycerol density gradient were diluted 1:1 with spraying buffer (200 mM ammonium acetate and 55% (v/v) glycerol) to a concentration of approximately 0.5 µM. Low-angle metal shadowing and electron microscopy was performed as described previously ^73^. In brief, 15 µL of sample was air-sprayed onto freshly cleaved mica pieces and then mounted and dried in a MED020 high-vacuum metal coater (Bal-tec). A Platinum layer of approximately 1 nm and a 7 nm carbon support layer were evaporated onto the rotating specimen at angles of 7 and 45 degrees, respectively. The Pt/C replicas were released from the mica on water, captured by freshly glow-discharged 400-mesh Pd/Cu or Cu grids (Plano GmbH), and visualized by transmission electron microscope (TEM). TEM was performed with a Morgagni 268D system (FEI) operated at 80 kV. Images were recorded on a 1376 x 1032 MegaView III CCD camera (Olympus) at nominal magnifications of 71.000, 89.000, and 110.000 resulting in pixel sizes of 0.856, 0.695, and 0.570 nm/pixel respectively and with a Tecnai T20 system (Thermo Scientific) operated at 200 kV and equipped with a 4096 x 4096 BM Eagle CCD camera (FEI) at nominal magnifications of 50.000, 62.000, and 80.000 resulting in pixel sizes of 0.2211, 0.1787, and 0.1382 nm/pixel respectively. Particles were manually selected and distances from CIP2A globular domains until the tail (dimers) or the end of the neighboring globular domain (tetramers) were measured using Fiji^74^. Data analysis and visualization was performed using matplotlib^75^.

### TOPBP1-CIP2A pulldown assays

A pGEX6P1 vector to express TOPBP1^32–1522^ was obtained from addgene (20375)^76^ and modified to express TOPBP1^549–996^. A CIP2A-binding deficient form was created by introducing eight mutations: V778A, L781A, F786A, V794A, F826A, L827A, F832A, and F836A (corresponding to residues in full-length protein). The generated TOPBP1^549–996^ constructs were transformed into *E. coli* BL21-CodonPlus(DE3)-RIL for protein expression. Cultures were inoculated from single colonies and grown at 37°C in LB containing Ampicillin and Chloramphenicol. At an OD_600_ of 0.6, expression was induced with 300 µM IPTG. Cells were kept at 16°C over-night for protein expression and harvested by centrifugation (3,000 *g*, 10 minutes, 4°C). After a wash with cold PBS, cell pellets from 500 mL cultures were flash-frozen in liquid nitrogen and stored at −70°C. All following steps were performed on ice or at 4°C. Cell pellets were resuspended in 25 mL lysis buffer (500 mM NaCl, 50 mM Tris-HCl pH 6.8, 2% (v/v) Glycerol, 10 mM BME, 10 mM Imidazole, 0.1% (v/v) Tween-20, 1 mM PMSF, 2 mM EDTA, 0.28 mg/mL Lysozyme). Cells were lysed by sonication, and the lysate was centrifuged at 20.000 *g*. Soluble material was incubated with approximately 1 mL equilibrated Ni-NTA agarose beads (Protino, Machery-Nagel) for at least 1 hour. Beads were pelleted by centrifugation at 500 *g* for 5 min and washed with wash buffer (500 mM NaCl, 50 mM Tris-HCl pH 6.8, 2% glycerol (v/v), 1 mM BME, 20 mM Imidazole) twice. For elution, beads were resuspended in 1 mL of buffer containing 320 mM Imidazole. Beads were pelleted by centrifugation and the supernatant containing GST-TOPBP1^549–996^–His was loaded onto columns for preparative size exclusion chromatography (SEC) with a Superdex 200 Increase 10/300 column (Cytiva) equilibrated in SEC buffer (50 mM Tris-HCl pH 6.8, 2% glycerol (v/v), 500 mM NaCl, 1 mM TCEP). Protein elution was followed using absorbance at 280 nm. Selected fractions were analysed by SDS-PAGE and Coomassie staining, snap frozen in liquid nitrogen, and stored at −70°C.

All steps of the pulldown assay were done on ice or at 4°C. Wild-type or mutant GST-TOPBP1^549–996^–His was incubated with approximately 10 µL equilibrated glutathione agarose beads (Protino, Machery-Nagel) in 1 mL binding buffer (500 mM NaCl, 50 mM Tris-HCl pH 6.8, 2% (v/v) glycerol) for at least 3 hours. Beads were pelleted by centrifugation at 600 *g* for 5 min. After binding of TOPBP1, the beads were washed with binding buffer twice. The washed beads were incubated with 100 nM wild-type or mutant full-length CIP2A in 500 µL binding buffer for 1 hour. The beads were washed four times with 1 mL of wash buffer (500 mM NaCl, 50 mM Tris-HCl pH 6.8, 2% (v/v) glycerol, 0.1% Tween20). Every wash included rotating the beads for 10 minutes. Before the 3^rd^ wash, the beads were transferred to a new tube. After the 4^th^ wash, the beads were pelleted, the supernatant was removed and the beads were resuspended and boiled in loading buffer for SDS-PAGE. 10 µL of the sample were loaded for SDS-PAGE and gels were either stained with Coomassie or used for Western blotting. CIP2A was detected using a polyclonal antibody against CIP2A (Thermo Fisher #PA5-83469), diluted 1:000, and a secondary antibody against rabbit conjugated to HRP, diluted 1:10 000, for chemiluminescence detection.

### Xenopus CIP2A peptide-IP

Mitotic *Xenopus laevis* egg extract was produced as previously described^77^. C-terminal peptides comprising the last 29aa of human CIP2A were produced by Genosphere Biotechnologies with a N-terminal biotin-SGSG-linker and a C-terminal carboxyl group (-COOH): wt: biotin-SGSGELNKHSHMIAMIHSLSGGKINPETVNLSI-COOH; SCR (scrambled): biotin-SGSGEIKNSGLENLVMGHHKSISSANMTIPLH-COOH; m4: biotin-SGSGELNKHSHENAMIHSRAGGKINPETVNLSI-COOH; m6: biotin-SGSGELNKHSHENAKIQSRAGGKINPETVNLSI-COOH. Lyophilized peptides were reconstituted in water and for each reaction 2 µg of peptide was incubated for 1 hour with 20 ul pre-calibrated streptavidin beads (Dynabeads MyOne Streptavidin C1, Invitrogen). Peptide-beads were washed three times with ELB-sucrose buffer, reconstituted in 10ul ELB-buffer (IP buffer (ELB-sucrose buffer: 10 mM HEPES-KOH ph7.7, 50 mM KCl, 2.5 mM MgCl_2_, 250 mM sucrose; 0.25 mg/mL BSA; 0.02% Tween-20), and incubated with 14 µl of protein extract for 30 min at room temperature. Subsequently, peptide-beads were washed by magnetic separation two times with 400 µl of IP Buffer (ELB-sucrose buffer; 0.25 mg/mL BSA; 0.02% Tween-20), two times with IP-buffer minus BSA, and lastly one time with ELB-sucrose buffer. Lastly, peptide-beads were resuspended in 2x Sample Buffer and proteins were eluded from peptide-beads by 5 min incubation at 100 °C.

### Cell culture

Human hTERT immortalized retinal pigmented epithelial RPE1 cells (CRL-4000) and human embryonic kidney HEK293T (CRL-3216) were obtained via the American Type Culture Collection (ATCC). RPE1 *TP53^−/−^ CIP2A^−/−^, TP53^−/−^ CIP2A^−/−^* CIP2A^wt^-V5, *CIP2A^−/−^* CIP2A^S904A^-V5, and *CIP2A^−/−^* CIP2A^ΔC^-V5 cells were described previously^23^. DLD1 wildtype and DLD1 *BRCA2^−/−^*human colorectal adenocarcinoma cells were obtained from Horizon (Cambridge, UK). RPE1 and HEK293T cells were cultured in Dulbecco’s modified Eagle medium (DMEM, Thermo Fisher) and DLD1, PEO1 and PEO4 cells were cultured in Roswell Park Memorial Institute (RPMI) medium. DMEM and RPMI were complemented with 10% (v/v) foetal calf serum (FCS), 1% penicillin and 1% streptomycin (Gibco). SUM149PT cells were cultured in Ham’s F12 Nutrient Mix, complemented with 5% (v/v) FCS, 1% penicillin, 1% streptomycin (Gibco), 10 μg/mL insulin (Lonza BioScience), 1 μg/mL hydrocortisone (Sigma). All cells with doxycycline-inducible CIP2A^wt^ and CIP2A^ΔC^ constructs were cultured in medium with Tet-System Approved FCS. RPE1, DLD1 and HEK293T cells were grown in a humidified incubator at 37° C, 5% CO_2_ and 20% O_2_. PEO1, PEO4, SUM149PT cells were grown in a humified incubator at 37°C, 5% CO_2_ and 3% O_2._ All cell lines were tested negative for *Mycoplasma* contamination.

### DNA cloning and mutagenesis

RPE1 *TP53^−/−^* cells harboring a *TP53* mutation in exon 4, and RPE1 *TP53^−/−^ CIP2A^−/−^* cells harboring a *CIP2A* mutation in exon 3 were previously described (*CIP2A^−/−^*cl#1, unless otherwise stated) ^78,79,23^. To generate RPE1 *TP53^−/−^ POLQ^−/−^*, a sgRNA (5’-GCCGGGCGGCGGGCTCAGCA–3’) targeting exon 1 was cloned in pSpCas9(BB)-2A-GFP (px458), kindly provided by Feng Zhang (Addgene #48138)^77,80^. Cells were transiently transfected using Fugene HD, and GFP-positive cells were subsequently sorted using a Sony Sorter SH800S, into monoclonal lines at 48 h after transfection. Mutations in *POLQ* were confirmed with Sanger sequencing (*POLQ^−/−^*: +1 insertion). SUM149PT parental and *BRCA1* reverted cells were previously described^81^. To generate SUM149PT *BRCA1^−/−^* clones, a sgRNA (5’–GAATGGATGGTACAGCTGTG– 3’) targeting exon 22 of *BRCA1* and was cloned in lentiCRISPRv2 plasmid (kindly provided by Feng Zhang, Addgene 52961). Cells were transfected using Xfect (Takara) and were subsequently selected using 2 ug/mL blasticidin at 24 hours after transfection. After selection, cells were plated in 96 wells to generate single clones and mutations in *BRCA1* were confirmed with Sanger sequencing (both clones: −1 and −2 deletion) and immunoblot analysis.

pMSCV CIP2A wt (1-905)-His, pMSCV CIP2A ΔC (1-876)-His, pMSCV CIP2A S904D-V5, pMSCV CIP2A S904E-V5, pTLCV2 CIP2A wt (1-905) and pTLCV2 CIP2A ΔC (1-876) were cloned using Gibson from previously reported pMSCV CIP2A wt (1-905)-V5 and pMSCV CIP2A-ΔC (1-876)-V5^23^. To generate doxycycline-inducible pTLCV2-CIP2A constructs, GFP-SLX4 was removed from pTLCV2 GFP-SLX4^82^ (kindly provided by Dr. Andrew Blackford) using restriction enzymes AgeI and NheI and replaced with either CIP2A wt or CIP2A ΔC constructs using Gibson cloning.

pLIB and pOPIN-B derived vectors were subsequently used to generate pMSCV CIP2A-His plasmids containing mutated CIP2A (m4, m6) and CIP2A chimeras via Gibson cloning. Of note, synonymous mutations in amino acids 442 and 443 (ACTGTC to ACCGTA) and amino acids 446 and 448 (ACCACT to ACAACC) were introduced in all pMSCV CIP2A-His plasmids to generate siRNA resistant cDNAs. pMSCV-CIP2A-His plasmids with m4, m6 and chimeric sequences consist of human CIP2A (1-869) followed by the following sequences: LVKLQQEELNKHSHENAMIHSR-AGGKINPETVNLSI (m4), LVKLQQEELNKHSHENAQIQSRAGGKINPETVNLSI (m6), QLEEIAQQLEEIAKQLKKIAE (synthetic), TISEKDRNIQRMKGELKELSRMRDMIFEL-TAKNKDDLNSS (*A. cephalotes*), QLQERKAQIESLEMEQKETEDLRKTIMSLMESKK-PKRKA (*D. melanogaster*), VIKEKTEELQEVKDNLAKHEQIAAMIHNLSSSKIPPKVSF (*O. bimaculoides*), DLDAARRNLQKLEQMREAMLSLASSSSN (*C. elegans*). These sequences are followed by these tags GLPETGGGHHHHTIESTNSTG (CIP2A-m4-His), GLPETGGGHHHHHH (all other CIP2A-His), GTKGNSADIQHSGGRSSLEGPR-FEGKPIPNPLLGLDSTRTGHHHHHH (CIP2A-V5-His).

### Viral transductions

For lentiviral transductions, HEK293T cells were transfected with pMD2.G and psPAX2 together with the indicated pTLCV2 plasmids. For retroviral transductions, HEK293T cells were transfected with pRetro-VSV-G, pRetro-gag/pol, pAdvantage with indicated pMSCV plasmids as described previously^12^. Virus-containing supernatant was harvested and filtered with a 0.45 μm syringe filter. Supernatant was supplemented with 4 μg/mL polybrene to infect target cells. Transduced cells were subsequently selected with 10 μg/mL blasticidin (pMSCV) or 1 μg/mL puromycin (pTLCV2). RPE1 *TP53^−/−^ CIP2A^−/−^* cells transduced with the various pMSCV CIP2A-His plasmids were sorted using a Sony SH800S Sorter to produce clonal lines. Clones were tested for levels of CIP2A-His expression levels, and clones with similar expression levels to CIP2A levels in parental RPE1 *TP53^−/−^* cells were selected for experiments. RPE1 *TP53^−/−^ CIP2A^−/−^* cells transduced with the pMSCV CIP2A-S904D and pMSCV CIP2A-S904E plasmids were checked via western blotting for similar expression levels to CIP2A-wt-V5 expressing cells.

### GFP-POLQ transfections

Indicated RPE1 cells were transfected with PiggyBac CAG eGFP-POLQ-Flag-P2A-Blast (kindly provided by dr. Raphael Ceccaldi)^28^ and pCMV-mPbase containing Piggybac transposase using FuGENE. GFP-positive cells were sorted at 48 h after transfection using a Sony SH800S Sorter and cultured in conditioned medium. 9-14 days after transfection, GFP-POLQ-expressing cells were sorted again using a Sony SH800S Sorter to yield polyclonal cell lines with low levels of GFP-POLQ expression.

### Microscopy

Immunofluorescence microscopy was performed as previously described^23^. In short, cells were seeded on glass coverslips in 12-well plates at 24-48 h prior to treatment. For micronuclei analysis, cells were left untreated or treated for 24 h with 200 nM aphidicolin (RPE1 cells), for 48 h with 0.1 μg/mL doxycycline (DLD1 cells) or 96 h with 1 μg/mL doxycycline (PEO1, PEO4 and SUM149PT cells).

For mitotic foci analysis, RPE1 cells were treated with 200 nM aphidicolin or 0.5 μM olaparib for 16 h. RPE1 cells were then treated with 5 μM RO-3306 for 4 h to delay G2-M transition. DLD1 cells were either pre-treated for 44 h with doxycycline prior to, or 4 h treated with doxycycline in combination with, 4 h 5 μM RO-3306 treatment. RO-3306 was washed out to promote progression into mitosis and cells were fixed after 30 min. with 2% paraformaldehyde in PBS or 2% paraformaldehyde with 0.1% Triton-X in PBS for 15 min. Cells were permeabilized with 0.5% Triton-X in PBS for 10 min and subsequently blocked with 2% BSA, 0.05% Tween in PBS (blocking buffer). Cells with incubated with primary antibodies in blocking buffer overnight at 4° C. The following antibodies were used: mouse-anti-CIP2A (1:500, Santa Cruz, sc-80659), rabbit-anti-CIP2A (1:300, Invitrogen, PA5-83469), mouse-anti-TOPBP1 (1:500, Santa Cruz, sc-271043), rabbit-anti-TOPBP1 (1:400, Bethyl, A300-111A-M) mouse-anti γH2AX (1:400, Millipore, 05-636, JBW301), mouse-anti-BTBD12 (SLX4) (1:500, Abnova, H00084464-B01P), rabbit anti-V5-tag (1:1000, Cell signaling, 12302S), mouse-anti-RPA32/RPA2 (1:500, Abcam, ab2175). After extensive washing, secondary antibody incubation was performed 1-2 h on room temperature (RT) in blocking buffer. The following secondary antibodies were used: Alexa 488 donkey-anti-mouse (1:500, Thermo Fisher, A21202), 488 donkey-anti-rabbit (1:500, Thermo Fisher, A21206), Alexa 647 donkey-anti-mouse (1:500, Thermo Fisher, A31571), Alexa 647 donkey-anti-rabbit (1:500, Thermo Fisher, A31573). DAPI staining was either performed directly after fixation for micronuclei analysis or immediately following secondary antibody incubation. After extensive washing, coverslips were mounted using either Prolong Gold antifade reagents (Life Technologies) or Mowiol (Merck).

For widefield microscopy, images were acquired using a Nikon Eclipse Ti-E inverted microscope equipped with a Hamamatsu C11440-22CU digital camera and NIS-Elements software (Nikon) or a Zeiss Axio Imager 2 with the Zen 3.0 software (Zeiss) using either a 10x, 40x or 63x objective. For analysis of mitotic cells, images at multiple z planes were taken at 1 μm distance, and images were then stacked. Foci analysis was done manually based on maximum image projections of the Z projections. CIP2A foci intensity was quantified using the FIJI (ImageJ) macro ‘Foci-analyzer’ (available at https://github.com/BioImaging-NKI/Foci-analyzer). Staining intensities were normalized per cell to the mean foci intensity of untreated RPE1 *TP53^−/−^* cells (Suppl. Fig 10E) or aphidicolin-treated RPE1 *TP53^−/−^ CIP2A^−/−^* cells reconstituted with CIP2A-wt-His (Fig. 2C) per individual experiment. Cells without any foci were excluded from this analysis.

### Immunoblotting

Inducible CIP2A^wt^ or CIP2A^ΔC^ overexpression was verified after treatment with 0.1 μg/mL doxycycline (DLD1) or 1 μg/mL doxycycline (PEO1, PEO4, SUM149PT) for 48 h before harvesting of the cell pellets. Cells were lysed with MPER (Thermo Fisher), complemented with protease and phosphatase inhibitor (Thermo Fisher). Protein concentrations were measured using BCA Protein Assay Kit (Pierce, Thermo Fisher). To validate *BRCA1* expression in SUM149PT cells, nuclear extracts (NE-PER, Thermo Fisher) were immunoblotted. Proteins were subsequently separated using SDS-polyacrylamide gels and transferred to PVDF membrane (Immobilon). Membranes were blocked with 5% skimmed milk in TBS with 0.05% Tween (TBS-T). Primary antibody incubation of membranes was performed overnight at 4°C with the following antibodies: mouse anti-CIP2A (1:500, Santa Cruz, sc-80659, C-terminal antibody), rabbit-anti CIP2A (1:1000, Invitrogen, PA5-83469, N-terminal antibody), mouse anti-V5-Tag (1:1000, Invitrogen, R960-25), rabbit anti-POLQ (1:1000), mouse anti-BRCA2 (1:1000, Calbiochem, OP95), mouse anti-Actin (1:10.000, MP Biomedicals, 691000), HSP90a/b (1:10.000, Santa Cruz, sc-13119), rabbit anti-α-Tubulin (1:1000, Cell signaling, 2125), mouse anti-BRCA1 (1:250, Merck, MS110). The membranes were subsequently incubated with either the HRP-conjugated goat anti-rabbit (1:5000, DAKO) or HRP-conjugated rabbit anti-mouse (1:5000, DAKO) secondary antibodies for 1 h at RT. Images were generated with a ChemiDoc MP Imaging (Bio-Rad) after incubation with Lumi-Light or SuperSignal West Femto Sensitivity Substrate (Thermofisher).

### Mutational signatures

Bulk mutational outcome profiles in stable Cas9-expressing IB10 (IB10-Cas9) mES cells and stable Cas9-expressing RPE1-hTERT cells were generated previously^31^. Three *CIP2A*-targeting sgRNAs and an additional five non-targeting sgRNAs were taken along in the mouse-specific and human-specific mini-pool for candidate-testing and are presented together with *POLQ*-, *POLL-*, and *FANCD2*-targeting sgRNAs as positive controls for TMEJ, c-NHEJ and HR-deficient backgrounds, respectively. Briefly, a strategy was employed in which a bulk population of Cas9-expressing cells were first transduced with a library of sgRNAs at MOI 0.1-0.2 and selected with polybrene, followed by the targeting of the library backbone using one of three sgRNAs (sgTargeting-1, sgTargeting-2 or sgTargeting-3, each targeting the same unique region within the library backbone at a different location) to generate DSB repair outcome spectra. POLQ-deletions contain the three most common POLQ-dependent deletions as previously has been described^31^.

To generate mutational signatures in RPE1 cells, between 180,000 and 250,000 cells were seeded in 6-well plates. While still in suspension, cells were transfected with RNP-complexes consisting of Alt-R S.p. Cas9 Nuclease V3 (IDT, 1081059), Alt-R CRISPR-Cas9 tracrRNA (IDT, 1072533) and Alt-R CRISPR-Cas9 crRNA targeting exon 1 of the *HBB* locus (‘5-GTAACGGCAGACTTCTCCTC-3’) (IDT) using RNAiMAX (Thermo Fisher). Medium was refreshed the following day, and cells were harvested approximately 48 h after transfection.

Genomic DNA was isolated according to a previously reported protocol^83^. In short, cells were lysed in lysis buffer (200 mM NaCl, 1% SDS, 10 mM Tris-HCl Ph 7.5, 10 mM EDTA) supplemented with 0.4 mg/mL Proteinase K. Lysates were incubated at 55°C for 3-5 h and subsequently neutralized using saturated NaCl. Lysates were centrifuged at 13,000 rpm for 15 min before adding one volume of isopropanol to supernatants to precipitate and extract DNA. Pellets were washed with 70% ethanol, and centrifuged at 13,000 rpm for 5 min. After centrifugation, ethanol was removed and pellets were air-dried and subsequently resuspended in TE buffer.

For targeted sequencing of the *HBB* locus, samples were processed as previously described^83^. In brief, primers specific to the target region (HBB_For: GATGTGTATAAGAGACAGGGCAGAGCCATCTATTGCTTA, HBB_Rev: CGTGTGCTCTTCCGATCTTATTGGTCTCCTTAAACCTGTCTT) were designed. Amplification was performed using Phusion High-Fidelity DNA Polymerase (Thermo Fisher Scientific) with the following conditions: initial denaturation at 95 °C for 10 minutes, followed by 25 cycles of 95 °C for 10 seconds, 60 °C for 30 seconds, and 72 °C for 30 seconds, with a final extension at 72 °C for 5 minutes. PCR products were purified using AMPure XP magnetic beads (Beckman Coulter) at a 1.2X ratio and eluted in 20 µl MQ.

Flow-cell adaptors and barcodes were added by performing a second PCR using 3 µl of the purified PCR1 product, 0.3 µM of the p5 and p7 index primers, Phusion High-Fidelity DNA Polymerase and the following conditions: 95 °C for 10 min, 7 cycles of 95 °C for 10 s, 58 °C for 30 s and 72 °C for 30 s, and the final extension 72 °C for 5 min. The PCR products were purified using AMPure XP beads at a 0.8X ratio and eluted in 20 µl MQ. The yield of each PCR sample was measured using the Quant-iT dsDNA assay kit (Thermo Fisher Scientific) and a iD3 microplate reader (SpectraMax), according to the manufacturer’s protocol. Samples were pooled at equimolar concentrations, and the final library was sequenced by 150-bp paired-end sequencing on a NovaSeq 6000 or NovaSeq X (Illumina). Next-generation sequencing data were processed using a custom JAVA program (SIQ), with the resulting output analysed and visualized using the SIQplotteR web tool (<u>siq.researchlumc.nl/SIQPlotteR/</u>)^84^. To normalize outcomes to the total number of mutation-containing reads, all unaltered (wildtype, wt) reads and SNVs were removed from the analysis. Only samples with more than 200,000 reads, including unaltered and SNVs, were included in our analysis.

### Clonogenic survival assays

DLD1 cells were plated at 100 cells/well (wt) or 300 cells/well (*BRCA2^−/−^*) in 6-well plates, in the absence or presence of 0.1 μg/mL doxycycline and fixed 10-14 days after seeding using 50% methanol, 29,95% water, 20% acetic acid and 0.05% Coomassie Brilliant Blue. Colonies were imaged using an EliSpot reader (Alpha Diagnostics International) with vSpot Spectrum software and manually counted. Approximately 2,500 PEO1, PEO4 or SUM149PT (parental, *BRCA1*-reverted, and *BRCA1^−/−^* cl#1 and cl#2) cells were plated at 2,500 cells/well in 12 well plates in the absence or presence of 1 μg/mL doxycycline. Medium was replaced including fresh doxycycline at 9 days after seeding. At 12 days after seeding, cellular viability was determined using CellTiter-Blue (Promega). Cells were subsequently fixed with 3.7% formaldehyde and stained with a 0,1% crystal violet solution. All clonogenic survival assays were seeded with three technical replicates per experiment. For DLD1 three biologically independent replicates were performed. For PEO1, PEO4 and SUM149PT three biological replicates were seeded in parallel.

### Generation of POLQ antibody

The POLQ antibody was raised in a rabbit that was immunized 5 times with protein fragment containing amino acids 1000–1350 of the human POLQ. This fragment was obtained by cloning into pET30a (Novagen) the corresponding cDNA and expressed *E. coli* and purified using Ni-NTA HisPur™ (Thermo Fisher) following the manufacturer’s instructions before injection. For affinity purification of the serum, approximately 50-100 μg of the POLQ antigen were loaded onto SDS–polyacrylamide gel electrophoresis and then transferred to a nitrocellulose membrane (Amersham Protran Premium 0.45, Cytiva) before staining with Ponceau S. The area of the membrane which contained the antigen was cut out, blocked with 2% BSA in TBS-T for 1 h and then incubated with the serum overnight at 4 °C. After 3 washes with TBS-T containing 0.5 M NaCl, bound antibodies were eluted with 100 M glycine-HCl, pH 2.3. After repeating this elution 3 times, the samples were pooled and 1 M Tris-HCl, pH 8.8, was immediately added to neutralize the pH of the antibody solution to pH 7.5.

### Nematode strains, culturing, and viability assays

All strains were derived from N2 Bristol unless otherwise stated and were cultivated under normal conditions^85^. Strains were generated using CRISPR/Cas9, following a published protocol^86^. A full list of strains used in this study is provided in Supplementary Table 1. The crRNAs, repair templates and genotypes used in this study can be found in Supplementary Table 2. Single worms (24 hours after L4 stage) were selected to NGM plates seeded with *E. coli* for 8 hours. Worms were recovered and transferred to NGM plates seeded with *E. coli*. Each day, worms were transferred to fresh plates for a total of 3 days. Eggs and the number of hatched worms were counted. The percentage viability was calculated as the number of hatched eggs/total number of laid eggs) × 100. The number of worms used per condition is indicated in the corresponding data set.

### Statistics and reproducibility

Numbers of measurement, number of replicates and explanations of mean, median and error bars are provided in the figure legends. Statistical tests were performed using GraphPad Prism 10 (GraphPad Software). All statistical tests on foci number and foci intensity were performed on the median per experiment.

### Data reporting

No statistical methods were used to predetermine sample sizes. The investigators were not blinded to allocation during experiments and outcome assessment.

## Data availability

All data from this study are available by request by contacting MATMvV or PJH. Sequencing data have been deposited into the NCBI SRA database. Source data will be provided with this paper.

**Generative AI** was not used for the preparation of the text in this manuscript.

---

**Supplementary Table 1.**
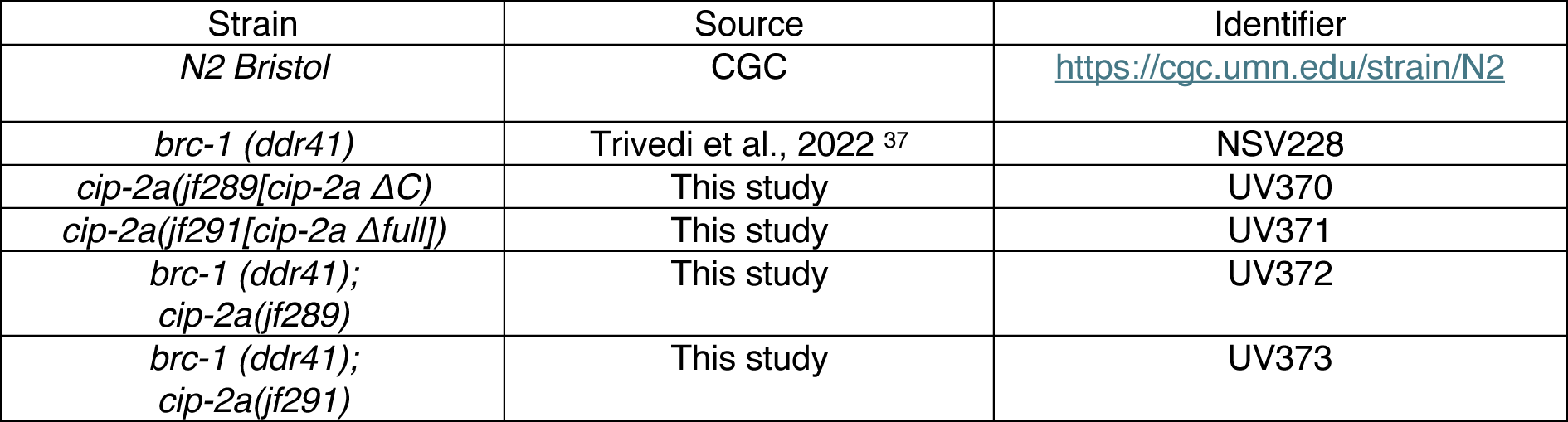

**Supplementary Table 2.**
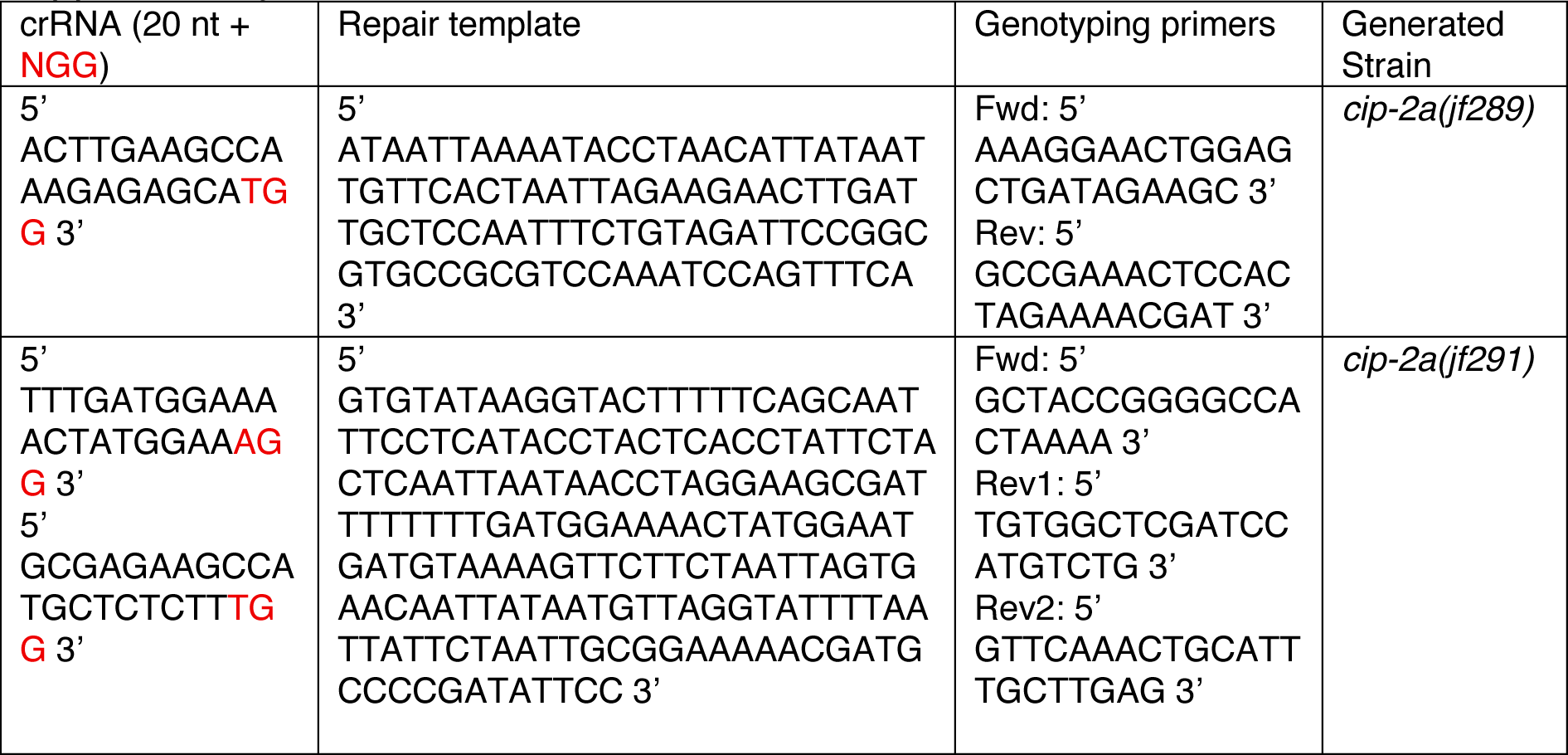

**Supplementary Figure 1.**
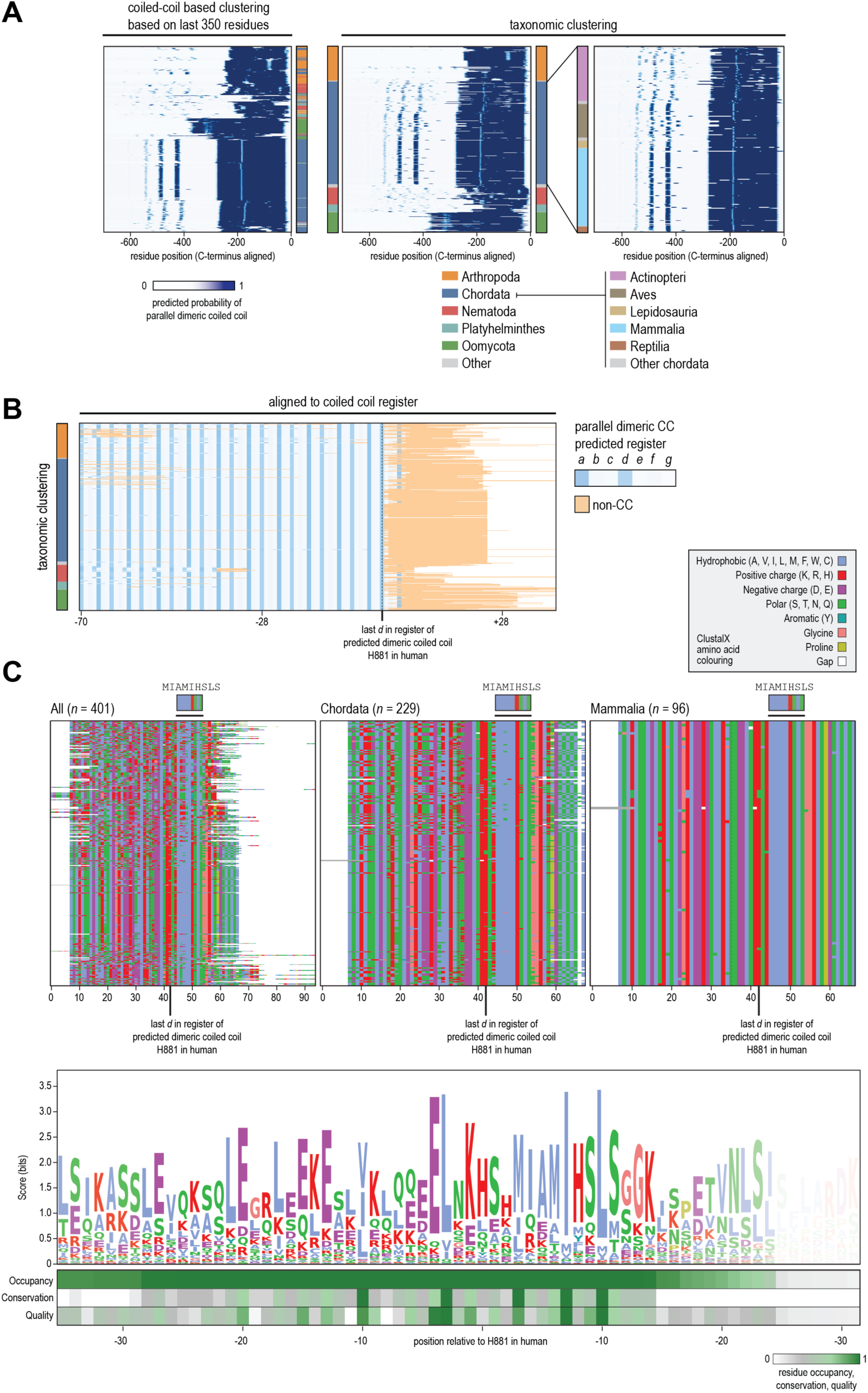
(**A**) The last 700 residues of 434 CIP2A homologs were scored for their tendency to form a dimeric parallel coiled coil (CC). All sequences were aligned with their ultimate residue on the right and clustered according to the CC pattern in the last 350 residues (left panel), or clustered taxonomically (middle panel). Most sequences (239) come from species belonging to the phylum chordata. The right panel details the classes with more than 5 entries found within chordata. (**B**) Sequences were aligned based on the predicted register of the parallel dimeric CC. Orange residues indicate non-CC sequences. Clustering according to taxonomy as in panel A. (**C**) CIP2A homologs with extensions of the parallel CC of <51 residues (401 in total), were aligned with a very high gap-penalty and analysed by colouring all residues according to their side-chain properties (left panel). A sequence logo was generated from this alignment, with the transparency per residue representing the alignment occupancy (see Fig 1F). In a similar manner, homologs from chordata (middle panel) and mammalia (right pannel) are shown. The conserved stretch at the end of the predicted parallel CC (in human MIAMIHSLS). The legend shows the ClustalX amino acid colouring scheme.

**Supplementary Figure 2.**
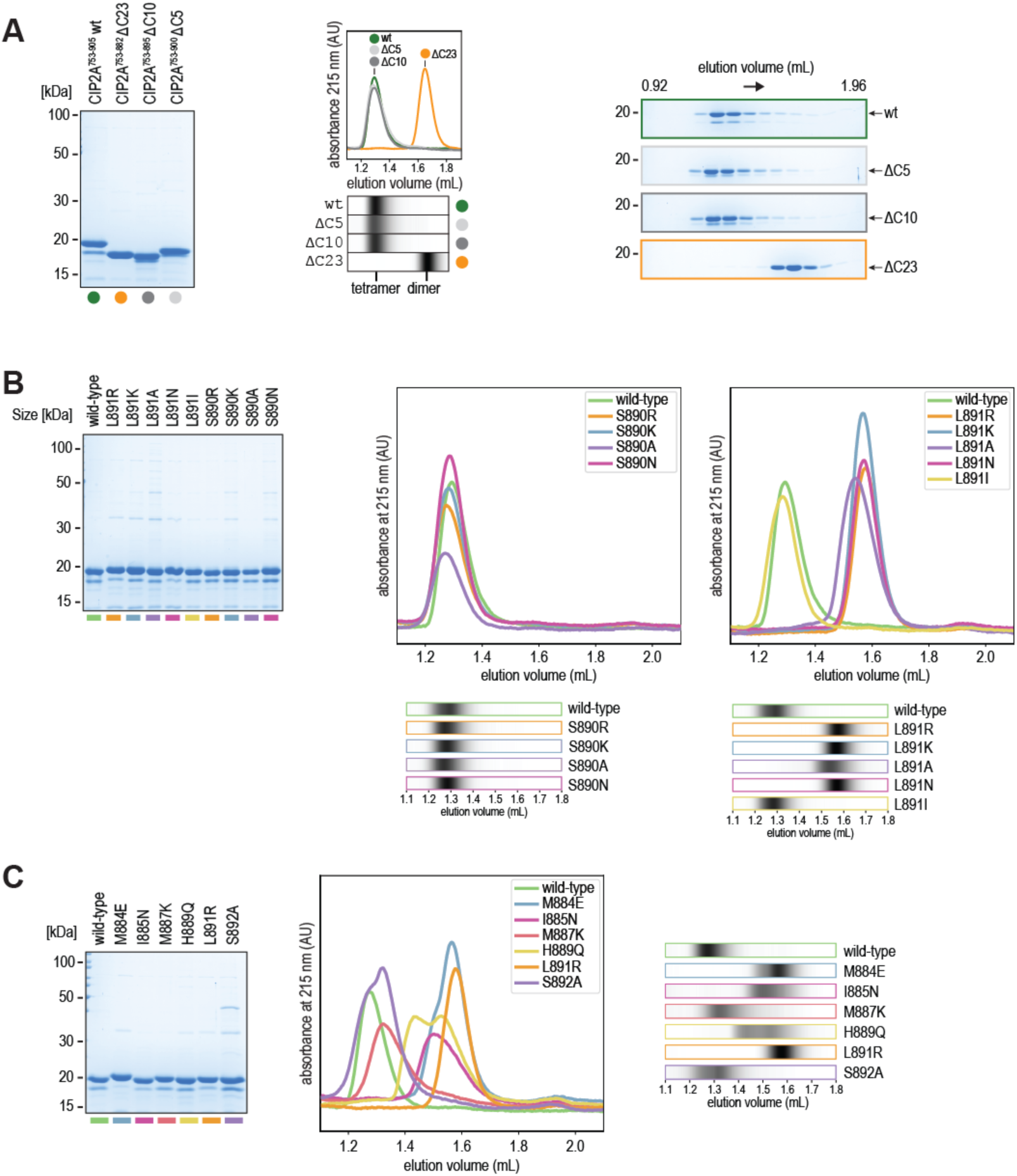
(**A**) Input samples for analytical size-exclusion chromatography (SEC) of CIP2A^753–905^ (wt), CIP2A^753–882^ (ΔC23), CIP2A^753–895^ (ΔC10) and CIP2A^753–900^ (ΔC5), analysed using SDS-PAGE and Coomassie staining (left) and the chromatograms from analytical SEC runs of these constructs depicted as traces and bar charts (middle, as in main Figure 1). For depiction of the chromatogram traces as bar charts, the absorbance of one sample was normalized from 0 to 1 within the depicted range of elution volume and subsequently adjusted according to the trace with the smallest area under the curve within one set. Coomassie-stained SDS-PAGE gels (right) show samples from each 80 µL SEC fraction. (**B**) Coomassie-stained SDS-PAGE gels of input samples for analytical SEC of CIP2A^753–905^ fragments with indicated single amino-acid mutations at position S890 and L891 (left). Original chromatogram profiles from analytical SEC runs of the different CIP2A fragments with representations as bar charts below (also shown in Fig. 1J). (**C**) Coomassie-stained SDS-PAGE gels of input samples for analytical SEC of CIP2A^753–905^ fragments with single amino-acid mutations at different positions (M884E, I885N, M887K, H889Q, L891R or S892A) (left). Original chromatogram profiles from SEC runs of the different CIP2A fragments with representations as bar charts below (also shown in Fig. 1J). Retention volumes indicate CIP2A tetramerization (1.2 – 1.4 mL) and dimerization (1.5 – 1.8 ml) in panels A-C.

**Supplementary Figure 3.**
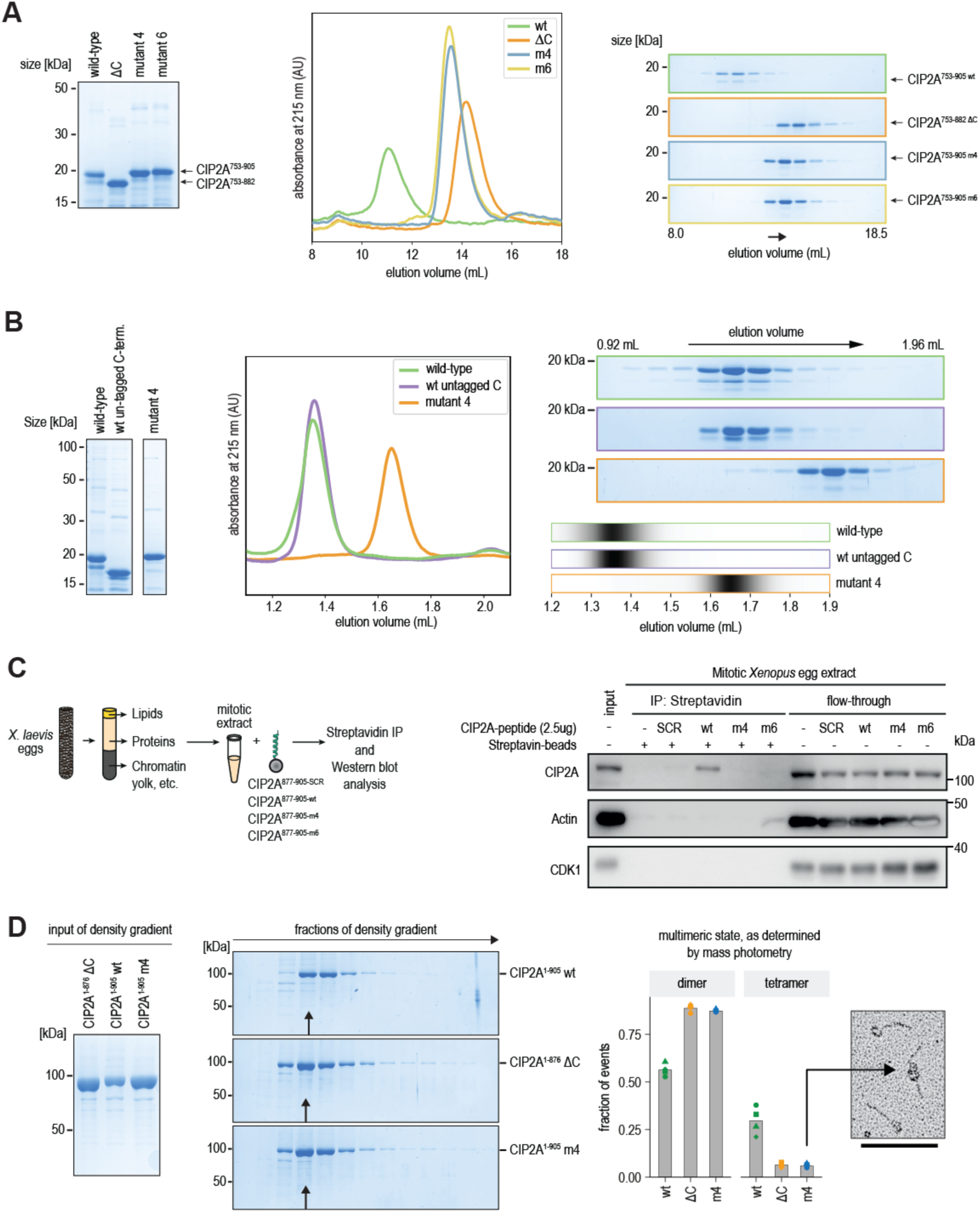
(**A**) Input samples for preparative size exclusion chromatography (SEC) of CIP2A wt, ΔC23, m4 (M884E, I885N, L891R, S892A) and m6 (m4 + M887Q, H889Q) fragments, analysed using SDS-PAGE and Coomassie staining (left). Chromatogram profiles of preparative SEC runs with CIP2A wt, ΔC23, m4 and m6 fragments (middle). Retention volumes indicate CIP2A tetramerization (10 – 12.5 mL) and dimerization (12.5 – 16 ml). Coomassie-stained SDS-PAGE gels of samples from each 750 µL fraction collected during preparative SEC (right). (**B**) Input samples for analytical SEC analysed by Coomassie-stained SDS-PAGE (left) and Chromatogram profiles from analytical SEC of CIP2A^753–905^(wt) and CIP2A^753–905^(mut4) with a C-terminal sortase-polyhistidine tag and CIP2A^753–905^(wt) without any C-terminal tag (middle). Coomassie-stained SDS-PAGE gels of samples from each 80 µL fraction collected during analytical SEC (top right). Chromatogram profiles shown in A depicted as bar charts (bottom right). (**C**) Mitotic *Xenopus Laevis* egg extracts were incubated with different biotinylated CIP2A^876–905^ peptides (wt, m4 and m6, which is m4 mutations plus M887K, H889Q) or a scrambled control peptide (CIP2A^SCR^), compromising the same 29 C-terminal amino acids in randomized order. Proteins bound to streptavidin beads and corresponding flow-through fractions were analysed by immunoblotting with indicated antibodies. (**D**) SDS-PAGE and Coomassie staining of Ni-NTA affinity purified full-length CIP2A wt, ΔC and m4 (M884E, I885N, L891R, S892A), which were used as input for sedimentation in a glycerol-based density gradient (left). Density gradient fractions (350 µL) were analysed by SDS-PAGE and Coomassie staining (right). The fractions chosen for further experiments are indicated by an arrow. The propensity of purified CIP2A wt, ΔC and m4 to form dimers or tetramers was measured using mass photometry. Examples of individual measurements are depicted in Fig. 1L. The fraction of events that can be attributed to dimers or tetramers was determined using Gaussian kernel density estimation in windows of 100 to 300 kDa and 300 to 500 kDa respectively. Symbols indicate technical replicates. Each 1-minute measurement captured between 6,383 and 14,958 events. Example of an electron micrograph of CIP2A^m4^ particles. Since end-end measurements were initiated on a globular domain (until the C-terminal end or until the end of the next globular domain), the shown particles would be measured as 4 dimers. However, the arrow indicates an apparent association between globular domains. This might explain the observed tetramers in CIP2A^m4^ and CIP2A^ΔC^ via mass photometry.

**Supplementary Figure 4.**
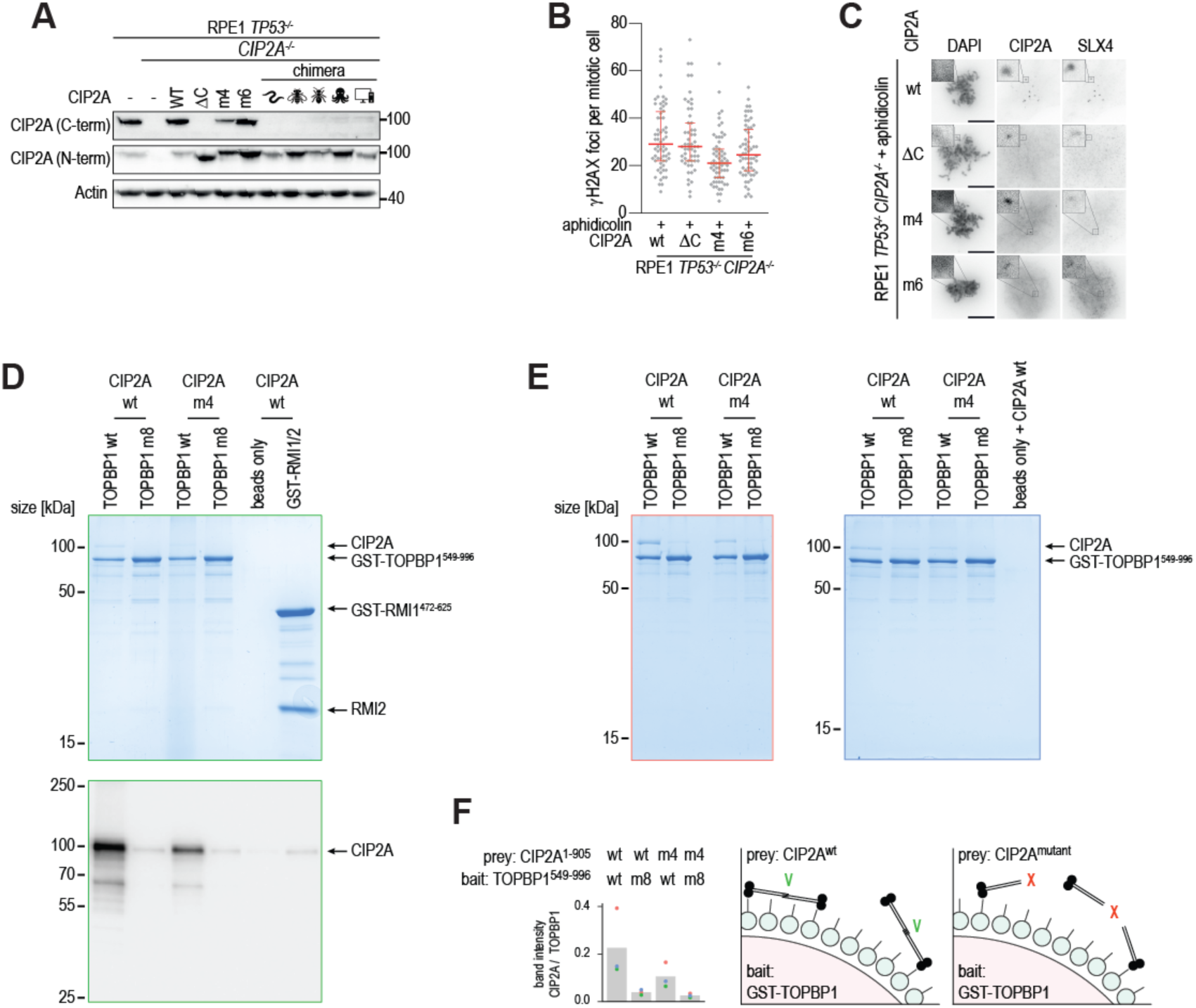
(**A**) Immunoblot analysis of RPE1 *TP53^−/−^* cells, RPE1 *TP53^−/−^ CIP2A^−/−^* cells or RPE1 *TP53^−/−^ CIP2A^−/−^* cells reconstituted with indicated with CIP2A variants. Lysates were immunoblotted for indicated proteins. (**B**) Quantification of the number of γH2AX foci per mitotic cell for *CIP2A^−/−^* cells reconstituted with indicated with CIP2A variants treated with aphidicolin (200 nM). Bars represent the median and interquartile range of two biological independent experiments with at least a total number of 55 cells measured per condition across experiments. (**C**) Representative wide-field images of RPE1 *TP53^−/−^ CIP2A^−/−^* cells reconstituted with indicated CIP2A variants stained for DAPI, CIP2A and SLX4 after aphidicolin treatment (200 nM). **(D)** Coomassie-stained SDS-PAGE gel with samples from a pull-down assay using purified TOPBP1^549–996^ as bait and purified CIP2A^1–905^ as prey (top) and western blot of CIP2A from the same samples (bottom). **(E)** Two additional replicates of the TOPBP1-CIP2A pull-down assay were analysed using SDS-PAGE and Coomassie staining. **(F)** Quantification of the three Coomassie-stained gels depicted in A and B. CIP2A band intensity was normalized to the TOPBP1 band intensity (left). Schematic representation of the CIP2A-TOPBP1 pull-down assay (right).

**Supplementary Figure 5.**
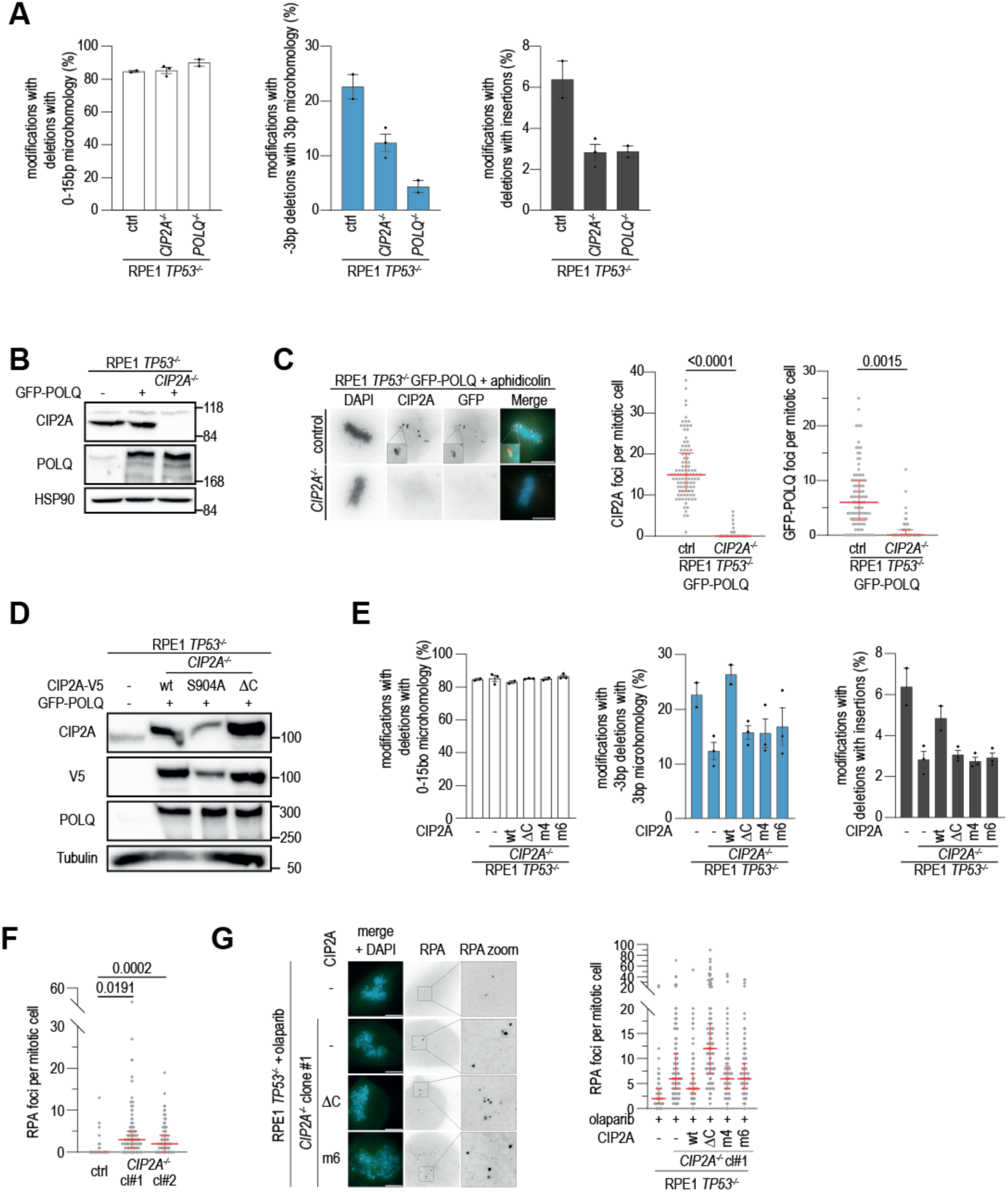
(**A**) Percentages of altered reads with deletions with 0-15 bp microhomology, 3bp deletions with 3bp microhomology and deletions with insertions are indicated for indicated RPE1 *TP53^−/−^* cells. Bars represent means and SEM of two (control, *POLQ^−/−^*) or three (*CIP2A^−/−^*) biologically independent experiments. (**B**) Immunoblot analysis of RPE1 *TP53^−/−^* cells and RPE1 *TP53^−/−^ CIP2A^−/−^* cells expressing GFP-POLQ. Lysates were immunoblotted for indicated proteins (**C**) Representative images of RPE1 *TP53^−/−^* and RPE1 *TP53^−/−^ CIP2A^−/−^* cells expressing GFP-POLQ stained for DAPI (blue), CIP2A (red) and GFP (green) after aphidicolin treatment (200 nM). Quantification of the number of CIP2A and GFP-POLQ foci per mitotic cell. Bars represent medians and interquartile range of three biologically independent experiments with at least 92 cells measured per condition across experiments. Two-tailed unpaired t-test was used on the medians per experiment. (**D**) Immunoblot analysis of RPE1 *TP53^−/−^*, and GFP-POLQ expressing *CIP2A^−/−^* cells, reconstituted with indicated CIP2A-V5 variants. Lysates were immunoblotted for indicated proteins. (**E**) Percentages of altered reads with deletions with 0-15 microhomology, 3bp deletions with 3bp microhomology and deletions with insertions are indicated for indicated RPE1 *TP53^−/−^* and *CIP2A^−/−^* cells reconstituted with indicated CIP2A variants. Bars represent means and SEM of two (control, wt) or three (*CIP2A^−/−^*, ΔC, m4, m6) biologically independent experiments. RPE1 *TP53^−/−^* and RPE1 *TP53^−/−^ CIP2A^−/−^* are identical to the samples plotted in panel A. (**F**) Quantification of RPA32 foci in RPE1 *TP53^−/−^* and indicated RPE1 *TP53^−/−^ CIP2A^−/−^* clones in unperturbed conditions. Bars represent the medians and interquartile range of three biologically independent experiments with at least 120 cells measured per condition across experiments. Two-tailed unpaired t-test was used on the medians per experiment. (**G**) Representative images of RPE1 *TP53^−/−^* cells, RPE1 *TP53^−/−^ CIP2A^−/−^* cells and RPE1 *TP53^−/−^ CIP2A^−/−^* cells reconstituted with indicated CIP2A variants treated with olaparib (0.5 μM) and stained for RPA32. Quantification of the number of RPA32 foci per mitotic cell. Bars for represent the median and interquartile range of three biologically independent experiments with >135 cells per condition measured across experiments. Scalebar throughout the figure represents 10 μm.

**Supplementary Figure 6.**
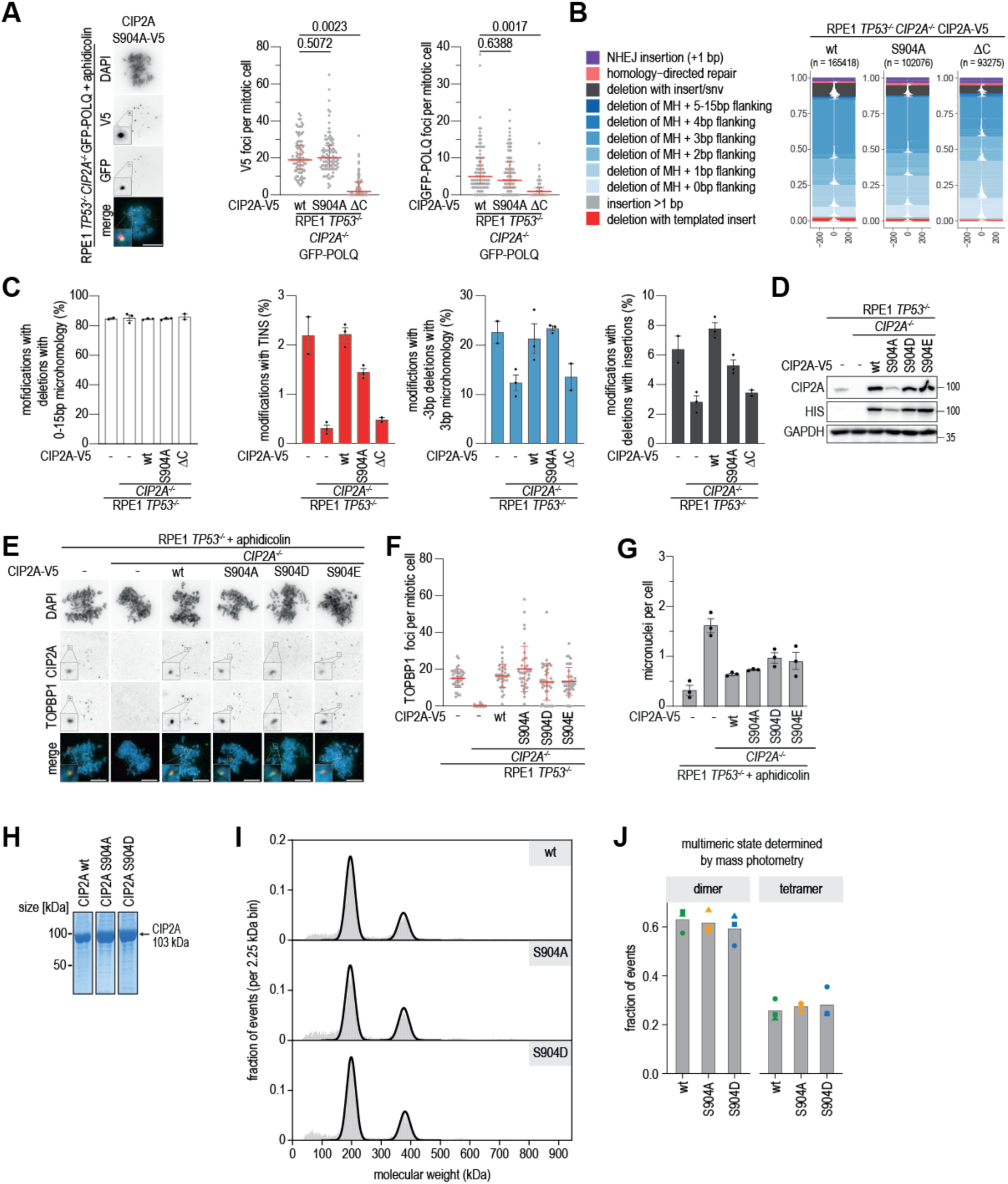
(**A**) Representative wide-field images of GFP-POLQ-expressing RPE1 *TP53^−/−^ CIP2A^−/−^* cells, reconstituted with indicated CIP2A-V5 variants treated with aphidicolin (200 nM). Quantification of the number of V5 foci and GFP-POLQ foci per mitotic cell. Bar represents the median and interquartile range of three biologically independent experiment with at least 94 cells measured per condition across experiments. Two-tailed unpaired t-test was used on the medians per experiment. Quantifications of GFP-POLQ foci of cells expressing CIP2A^wt^ and CIP2A^ΔC^ are identical to samples plotted in Fig. 2I. (**B**) Tornado plots of repair outcomes for *CIP2A^−/−^* cells reconstituted with indicated CIP2A-V5 variants. ‘n’ represents the number of reads plotted per tornado plot. X-axis indicates the distance relatively to the breaksite at 0. (**C**) Percentages of altered reads with deletions with 0-15 microhomology, deletions with TINS, 3bp deletions with 3bp microhomology and deletions with insertions are indicated for cells in panel B. Bars represent means and SEM of two (RPE1 *TP53^−/−^*, ΔC) or three (*CIP2A^−/−^*, wt, S904A) biologically independent experiments. RPE1 *TP53^−/−^* and *CIP2A^−/−^* are identical to the samples plotted in Fig. 2 and Supplementary Figure 4D. (**D**) Immunoblot analysis of RPE1 *TP53^−/−^* cells, *CIP2A^−/−^* cells or *CIP2A^−/−^* cells reconstituted with indicated with CIP2A-V5 variants. Lysates were immunoblotted for indicated proteins. (**E**) Representative wide-field images of RPE1 *TP53^−/−^*, *CIP2A^−/−^* and *CIP2A^−/−^* cells reconstituted with indicated CIP2A-V5 variants treated with aphidicolin (200 nM). (**F**) Quantification of the number of TOPBP1 foci per mitotic cell. Bar represents the mean and SD with at least a total of 30 cells measured per experimental condition. Scalebar represents 10 μm. (**G**) Quantification of micronuclei per cell in RPE1 *TP53^−/−^*, *CIP2A^−/−^* and *CIP2A^−/−^* cells reconstituted with indicated CIP2A-V5 variants treated with aphidicolin (200 nM). Bars represent means and SEM of four biologically independent experiments, with >845 cells per condition measured across experiments. **(H)** Coomassie-stained SDS-PAGE of samples from the density gradient fractions that were used for mass photometry measurements of CIP2A^1–905^ wild-type, S904A and S904D. **(I)** Representative mass photometry measurements of CIP2A^1–905^ wild-type, S904A and S904D. Bin size is 2.25 kDa. Black traces show Gaussian kernel density estimation (KDE) to obtain a bin-independent representation of mass distributions in the 100-300 kDa (dimer) and 300-500 kDa (tetramer) range. **(J)** Fraction of dimers and tetramers observed in three mass photometry measurements of CIP2A wild-type, S904A or S904D. Symbols indicate technical replicates. Per 1-minute measurement between 7,849 and 10,880 events were measured.

**Supplementary Figure 7.**
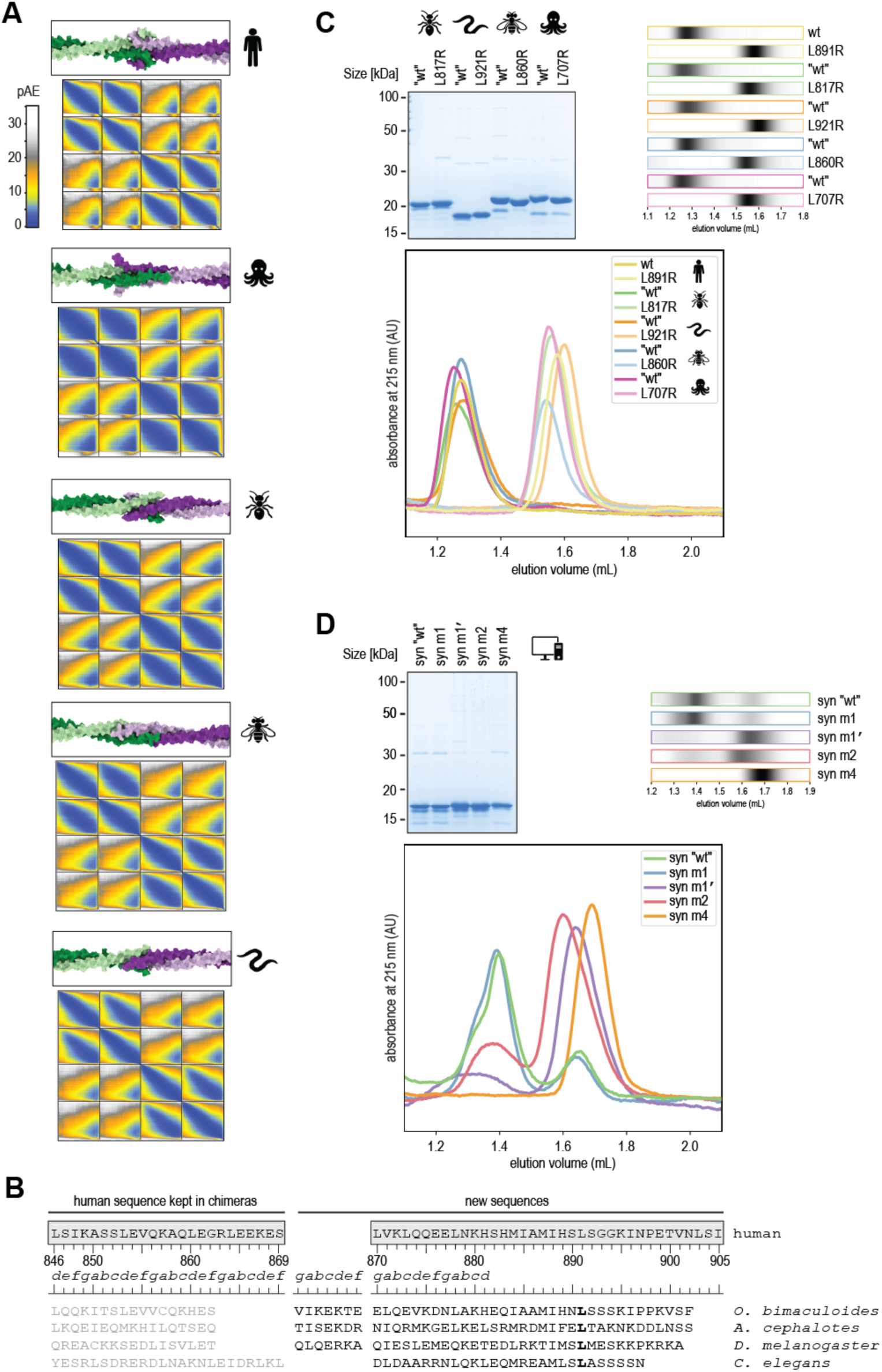
(**A**) Predicted Aligned Errors following Alphafold predictions of four copies the C-terminal, as shown in Fig. 3B. Low errors (dark blue) in the top right and bottom left quadrants indicate proximity between the C-termini of the dimers (green and the purple) for all species. (**B**) Detailed information on the design of chimeric CIP2A. Letters *a-g* indicate the predicted registered of the parallel dimeric coiled coil. (**C**) Affinity-purified CIP2A chimeric fragments were analysed using SDS-PAGE and Coomassie staining before size exclusion chromatography (SEC) (top left). For human CIP2A^wt^ and L981R fragments see Suppl. Fig. 2C. Chromatogram profiles from SEC runs of the CIP2A chimeric fragments and human CIP2A showing absorbance at 215 nm are depicted as original traces (bottom) and as grey scale bar charts (top right, also shown in Fig. 3E). For depiction of the traces as bar charts, the absorbance of one sample was normalized from 0 to 1 within the depicted range of elution volume and subsequently adjusted according to the trace with the smallest area under the curve within one set. **(D)** Coomassie-stained SDS-PAGE gels of affinity purified CIP2A constructs containing an intact synthetic C-terminal tetramerization domain or mutated synthetic tetramerization domain (top left). Original chromatogram traces from SEC runs of the different synthetic CIP2A fragments (bottom) with representations as bar charts on top (also shown in Fig. 3E)

**Supplementary Figure 8.**
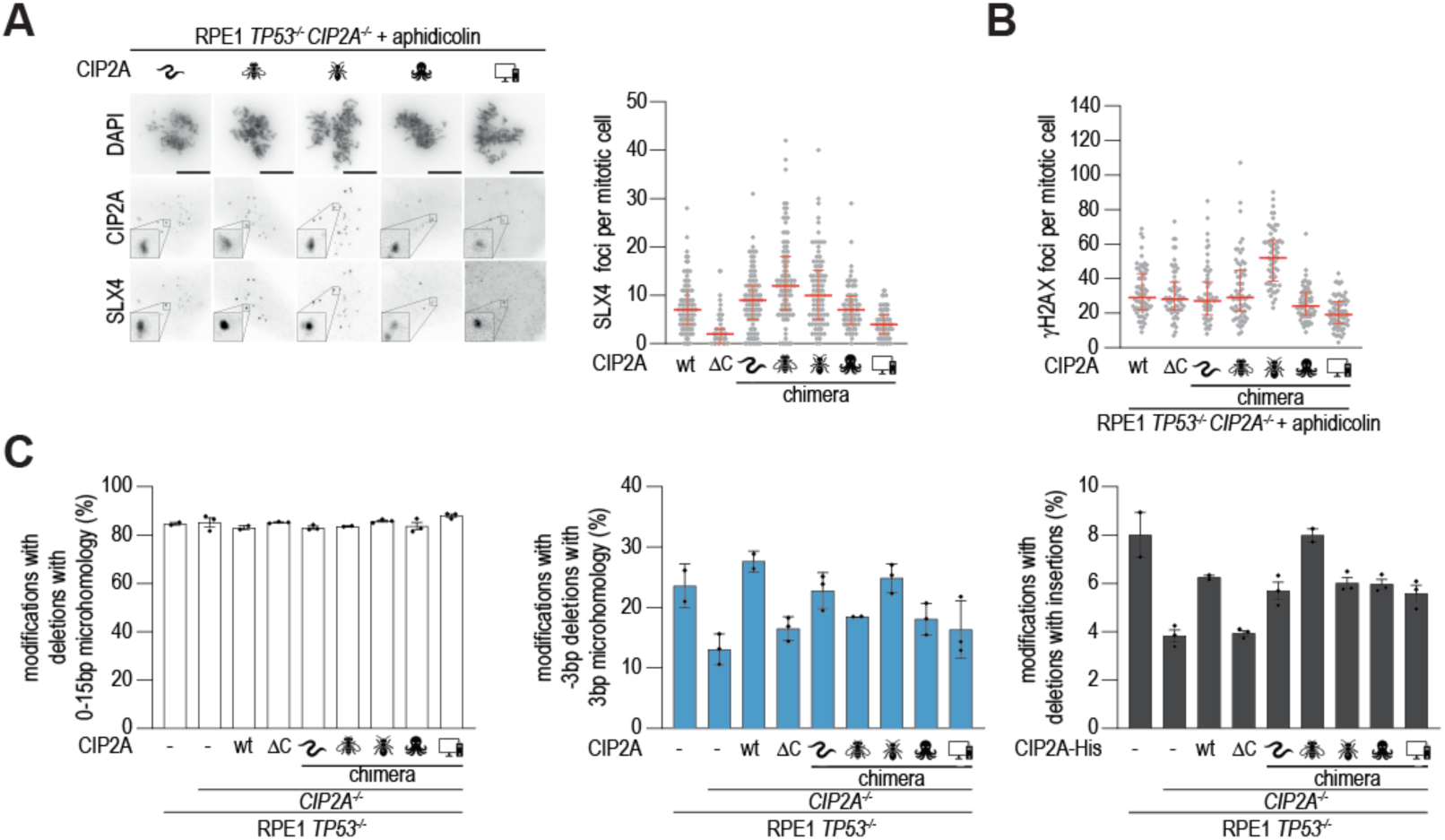
**(A)** Representative wide-field images of *CIP2A^−/−^* cells reconstituted with indicated CIP2A variants stained for DAPI, CIP2A and SLX4 treated with aphidicolin (200 nM). Scalebar: 10 μm. Quantification of the number of SLX4 foci per mitotic cell. Bars represent medians and interquartile range of three biological independent experiments with at least 90 cells measured per condition across experiments. (**B**) Quantification of the number of γH2AX foci per mitotic cell for *CIP2A^−/−^* cells reconstituted with indicated with CIP2A variants treated with aphidicolin (200 nM). Bars represent the median and interquartile range of two biologically independent experiments with at least a total number of 55 cells measured per condition across experiments. wt and ΔC are identical to the samples plotted in Suppl. Fig. 4B. (**C**) Percentages of altered reads with deletions with 0-15 microhomology, 3bp deletions with 3bp microhomology and deletions with insertions are indicated for *CIP2A^−/−^* cells reconstituted with indicated CIP2A variants. Bars represent the mean and SEM of two (RPE1 *TP53^−/−^*, wt and *D. melanogaster*) or three (*CIP2A^−/−^*, ΔC, *C. elegans*, *A. cephalotes*, *O. bimaculoides* and synthetic) biologically independent experiments. RPE1 *TP53^−/−^*, *CIP2A^−/−^*, wt and ΔC are identical to samples plotted in Fig. 2. ‘n’ represents the number of reads plotted per tornado plot.

**Supplementary Figure 9.**
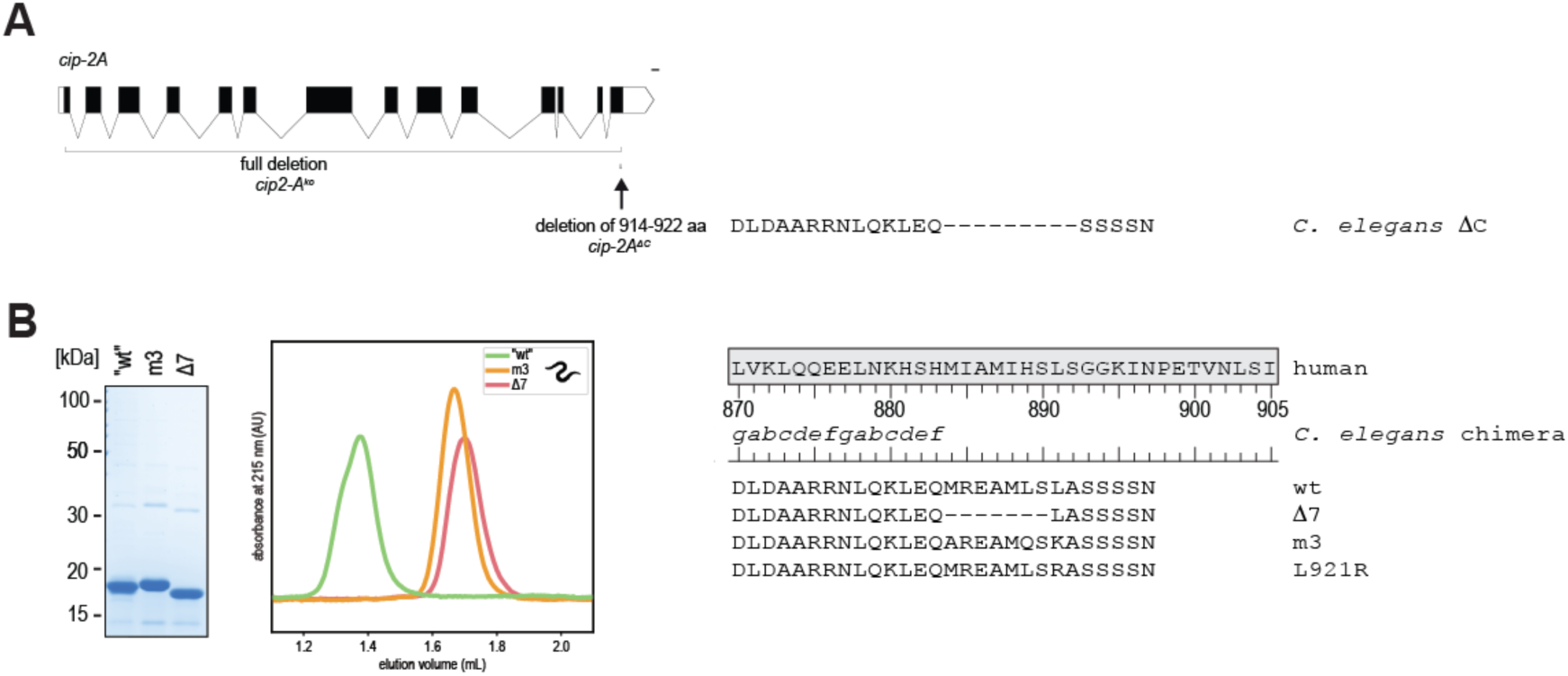
**A**) Schematic representation of the *C. elegans cip-2A* gene and the two generated alleles: *cip-2A* full deletion (*cip-2A^ko^*) and *cip-2A^ΔC^*. (**B**) The chimera protein consisting of human CIP2A753-869 and C. elegans CIP-2A900-927 (“wt”) and 2 mutated versions (m3 and Δ7) were analysed by size-exclusion chromatography as described in Figure 1H. The “wt” trace is also shown in Suppl. Figure 7C. The L921R mutant is shown in Figure 7C.

**Supplementary Figure 10.**
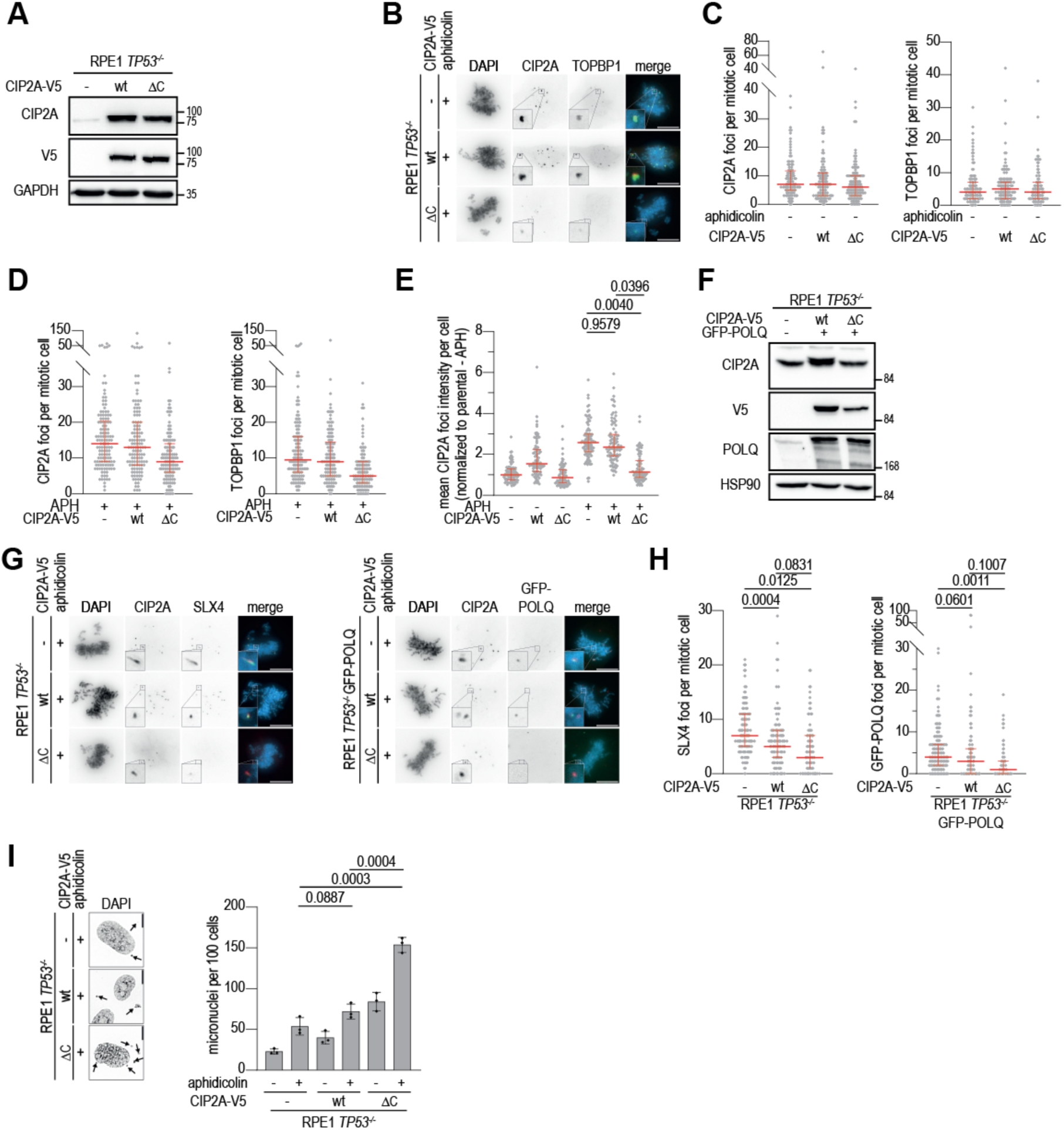
(**A**) Immunoblot analysis of RPE1 *TP53^−/−^* cells and RPE1 *TP53^−/−^* cells transduced with indicated CIP2A-V5 variants. Lysates were immunoblotted of indicated proteins. (**B**) Representative wide-field images of cells described in panel A stained for DAPI (blue), CIP2A (green) and TOPBP1 (red) after aphidicolin treatment (200 nM). (**C-E**) Quantification of the number of CIP2A and TOPBP1 foci and the mean CIP2A foci intensity per cell for cells either left untreated or treated with aphidicolin (200 nM), described in panel A. Bars represent medians and interquartile range of four biological replications with 123 cells (panel C, D) or three biologically replicates with 90 cells (panel E) measured per condition across experiments. Two-tailed unpaired t-test was used on the medians per experiment. (**F**) immunoblot analysis of RPE1 *TP53^−/−^* cells and RPE1 *TP53^−/−^* cells transduced with indicated CIP2A-V5 variants and stably transfected with GFP-POLQ. Lysates were immunoblotted for indicated proteins. (**G**) Representative wide-field images of cells described in panel A stained for DAPI (blue), CIP2A (red) and SLX4 (green), and representative wide-field images of cells described in panel G stained for DAPI (blue), CIP2A (red) and GFP (green) after aphidicolin treatment (200 nM). (**H**) Quantification of the number of SLX4 and GFP-POLQ foci. Bar represents the median and interquartile range of four biological independent experiments with at least 125 cells measured per condition across experiments. Two-tailed unpaired t-test was used on the medians per experiment. (**I**) Representative wide-field images of cells described in panel A stained for DAPI after aphidicolin treatment (200 nM). Arrows indicate micronuclei. Quantification of the number of micronuclei per 100 cells for untreated cells or cells treated with aphidicolin (200 nM). Bars represent the mean and SEM of three biologically independent experiments with at least a total of 417 cells measured per condition across experiments. Two-tailed unpaired was used. Scalebars throughout the figure represents 10 μm.

**Supplementary Figure 11.**
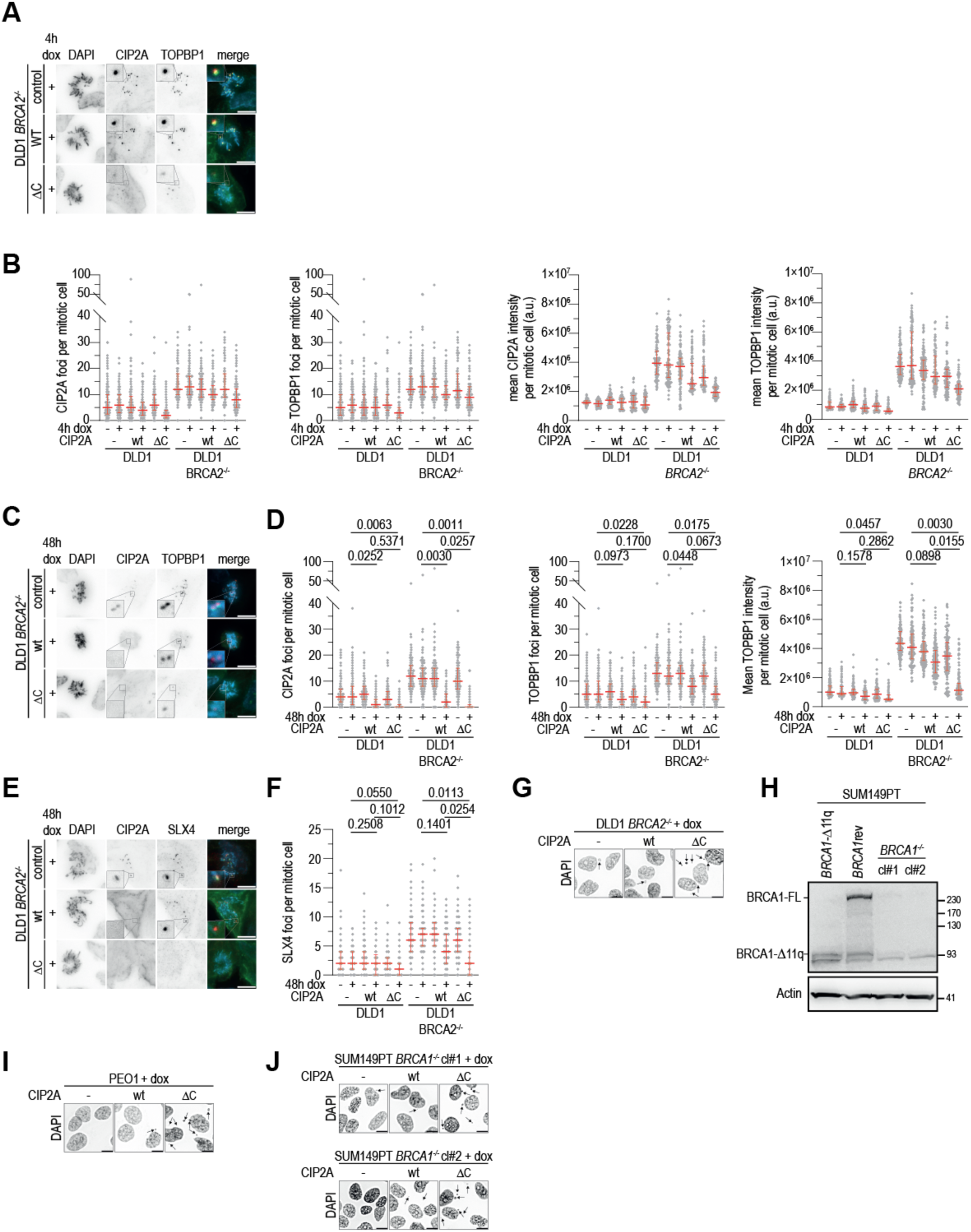
(**A, C**) Representative wide-field images of *BRCA2^−/−^* and *BRCA2^−/−^* DLD1 cells transduced with CIP2A^wt^ or CIP2A^ΔC^ treated for 4 hours or 48 hours with doxycycline (dox, 0.1 μg/mL) and stained for DAPI (blue), CIP2A (green) and TOPBP1 (red). (**B, D**) Quantification of the number of CIP2A and TOPBP1 foci, and mean foci intensity for conditions as described in panels A and C. Bars represent the median and interquartile range of three biologically independent experiments with at least a total of 94 (panel B) or 119 (panel D) measured cells per condition across experiments. Two-tailed unpaired t-test was used on the medians per experiment. (**E**) Parental *BRCA2^−/−^* DLD1 cells and *BRCA2^−/−^* DLD1 cells transduced with CIP2A^wt^ or CIP2A^ΔC^ were treated for 48 hours with dox (0.1 μg/mL) and stained for DAPI (blue), CIP2A (green) and SLX4 (red). (**F**) Quantification of the number of SLX4 foci per cell as described in panel E. Bars represent the median and interquartile range of three biologically independent experiments with at least 123 cells measured per condition across experiments. Two-tailed unpaired t-test was used on the medians per experiment. (**G)** Representative wide-field images of *BRCA2^−/−^* and *BRCA2^−/−^* DLD1 cells transduced with CIP2A^wt^ or CIP2A^ΔC^, treated with dox (0.1 μg/mL) and stained for DAPI. (**H**) Immunoblot analysis of parental (BRCA1 Δ11q), *BRCA1* reverted, *BRCA1^−/−^* cl#1 and *BRCA1^−/−^* cl#2 SUM194PT cells. Lysates were immunoblotted for indicated proteins. (**I, J**) Representative wide-field images of PEO1 cells and SUM149PT *BRCA1^−/−^* cl#1 and cl#2 cells transduced with CIP2A^wt^ or CIP2A^ΔC^, treated with dox (1 μg/mL) and stained for DAPI. Scalebars throughout the figure represent 10 μm, and arrows indicate micronuclei.

